# Phosphate starvation-induced CORNICHON HOMOLOG 5 as endoplasmic reticulum cargo receptor for PHT1 transporters in *Arabidopsis*

**DOI:** 10.1101/2024.06.20.599911

**Authors:** Chang-Yi Chiu, Cheng-Da Tsai, Hui-Fang Lung, Jhih-Yi Wang, Ming-Hsuan Tsai, Alastair J. McGinness, Satomi Kanno, Verena Kriechbaumer, Chiao-An Lu, Tzu-Yin Liu

## Abstract

Inorganic phosphate (Pi) is essential for plant growth and is acquired and distributed by the plasma membrane PHOSPHATE TRANSPORTER 1 proteins (PHT1s). Enhancing the abundance of PHT1s at the cell surface thus ensures plant productivity and sustainable agriculture. CORNICHON HOMOLOG proteins (CNIHs) are conserved eukaryotic cargo receptors that mediate the selective endoplasmic reticulum (ER) export of membrane proteins. In this study, we identified the *Arabidopsis thaliana CNIH5* (*AtCNIH5*) as a Pi starvation-inducible gene, preferentially expressed in vascular tissues and outer root cell layers above the meristem. *At*CNIH5 co-localizes to the *At*SAR1A/*At*SEC16A/*At*SEC24A-labeled ER exit sites and interacts with *At*PHT1;1. Loss of *AtCNIH5* confers reduced shoot Pi levels under Pi sufficiency due to the reduced translocation of Pi from roots to shoots, as well as decreased Pi uptake under Pi deficiency. The *cnih5* mutant exhibits decreased abundance of *At*PHT1s but increased PHOSPHATE TRANSPORTER TRAFFIC FACILITATOR1 (*At*PHF1), which enables the ER exit of PHT1s. The *cnih5* mutant also shows a lower plasma membrane targeting efficiency of split-GFP tagged*-At*PHT1;1 in the root hair and the epidermis within the root transition/elongation zone. Consistently, dysfunctional *At*CNIH5 exerts a suppressive effect on the growth of *phf1* and alleviates Pi toxicity in the Pi overaccumulator *pho2*. However, the *in vivo* protein–protein interaction and degradation assays indicated that *At*CNIH5 is not a direct target of *At*PHO2. Our findings unveil that *At*CNIH5 is a low Pi-responsive ER cargo receptor that interplays with *At*PHF1 to promote the plasma membrane targeting of *At*PHT1s in a cell-type-dependent manner.

## Introduction

Membrane trafficking that dictates the plasma membrane (PM) proteins in reaching their destination is intimately linked with plants’ plasticity to changing environments and, thus, crucial for plant growth and fitness under adverse conditions. Among the endoplasmic reticulum (ER) accessory proteins involved in membrane trafficking, CORNICHON HOMOLOG (CNIH) proteins are responsible for the selective export of membrane cargoes to the Golgi or post-Golgi compartments by cycling between the ER and the Golgi (Powers & Barlowe, 1998, Harmel *et al*., 2012). They are conserved in eukaryotes and presumably harbour three membrane-spanning α-helixes with the N-terminus towards the cytoplasm based on the “positive-inside rule” and supported by some experimental evidence (Powers & Barlowe, 2002). However, cryo-electron microscopy revealed that the N– and C-termini of mammalian CNIH2 and CNIH3 are located in the extracellular space (Nakagawa, 2019, Zhang *et al*., 2021, Gangwar *et al*., 2023). The AlphaFold model also predicts that the budding yeast *Saccharomyces cerevisiae Sc*CNIH/Erv14 has a topology of four transmembrane domains with both termini located in the ER lumen (Lagunas-Gomez *et al*., 2023).

The *Drosophila* cornichon is the first identified CNIH that promotes the ER export of the transforming growth factor α (TGFα)-like growth factor and its trafficking towards the oocyte surface during oogenesis (Roth *et al*., 1995, Bökel *et al*., 2006). In yeast, systematic genetic manipulations combined with automated microscopy screening uncovered that Erv14 serves as an ER cargo receptor of about 40 membrane cargoes with diverse functions, including proteins involved in transport, budding, sporulation, and cell wall synthesis (Powers & Barlowe, 2002, Nakanishi *et al*., 2007, Herzig *et al*., 2012, Pagant *et al*., 2015, Zimmermannová *et al*., 2019, Powers & Barlowe, 1998). Of note, yeast cells lacking Erv14 showed normal PM localization of the K^+^-importer Trk2 yet mistargeting its paralog Trk1, despite that Trk2 is nearly 55% identical to Trk1 (Zimmermannová *et al*., 2019), suggesting a regulatory mechanism operating in the cargo selectivity of CNIHs. For instance, Erv14 employs multiple modes of cargo binding to meet the different requirements of various cargo proteins. It simultaneously binds to the membrane cargo and the coat protein complex II (COPII) subunit Sec24, thus incorporating the membrane cargo into the COPII carriers (Powers & Barlowe, 1998, Powers & Barlowe, 2002, Pagant *et al*., 2015). Similarly, Sec24 was reported to recognize two signals to efficiently drive the ER exit of client proteins, one on membrane cargoes and the other on Erv14 (Pagant *et al*., 2015).

Despite plant CNIH proteins (CNIHs) being less functionally characterized, they seem to have a similar role to their yeast counterpart. The moss (*Physcomitrium paten*s) *Pp*CNIH2 was identified as a specific cargo receptor of the auxin efflux transporter homolog PINA for its polar localization in protonema cells (Yáñez-Domínguez *et al*., 2023). The rice (*Oryza sativa*) *Os*CNIH1 was shown to mediate the Golgi targeting of the sodium transporter *Os*HKT1;3 (Rosas-Santiago *et al*., 2015). Moreover, overexpression of the pumpkin (*Cucurbita moschata* var. *Rifu*) *Cm*CNIH1 improves the salt tolerance by regulating the PM localization of the sodium-selective transporter *Cm*HKT1;1 (Wei *et al*., 2023). The *Arabidopsis cnih1*, *cnih4*, and *cnih1/4* double mutants were shown to exhibit reduced Ca^2+^ fluxes in the pollen tube tip due to the impaired targeting of glutamate receptor-like channels (GLRs) (Wudick *et al*., 2018). Furthermore, the patch-clamp analysis of mammalian COS-7 cells expressing *At*GLR3.3 together with *At*CNIH1 and *At*CNIH4 displayed a current two-fold relative to those co-expressing *At*GLR3.3 with *At*CNIH1 or *At*CNIH4 alone, indicating that the formation of *At*CNIH oligomers potentially imposes an additional layer of regulation on cellular Ca^2+^ homeostasis (Wudick *et al*., 2018). Therefore, by modulating the trafficking of various transporters along the secretory pathway, the plant CNIH family can regulate various ion homeostasis. Phosphate (Pi) is known to be one of the most limiting mineral nutrients for plant growth and productivity. Due to its chemical properties, Pi is primarily immobilized in the soil, resulting in poor (Manning, 2008).bioavailability (Manning, 2008). To adapt to low Pi availability, plants employ a series of strategies, including enhancing Pi acquisition through the PM-localized PHOSPHATE TRANSPORTER 1 (PHT1) family (Poirier & Bucher, 2002, Młodzińska & Zboińska, 2016). In *Arabidopsis*, *At*PHT1;1 and *At*PHT1;4 are responsible for up to 75% of Pi uptake under a wide range of Pi concentrations and thus play dominant roles in Pi acquisition (Karthikeyan *et al*., 2002, Mudge *et al*., 2002, Shin *et al*., 2004, Ayadi *et al*., 2015, Misson *et al*., 2004). Under Pi sufficiency, *At*PHT1;1 also plays an important role in Pi translocation from roots to leaves (Ayadi *et al*., 2015). *At*PHT1;5 participates in Pi mobilization from source to sink organs (Nagarajan *et al*., 2011). *At*PHT1;8 and *At*PHT1;9 contribute to both Pi uptake and Pi translocation toward the vascular tissues in roots under Pi starvation (Remy *et al*., 2012, Lapis-Gaza *et al*., 2014).

Under Pi repletion, *AtPHT1;1* shows a robust gene expression in the root hair-producing trichoblast and is the most abundant member (Mudge *et al*., 2002, Karthikeyan *et al*., 2002, Ayadi *et al*., 2015). In response to low Pi, *AtPHT1;1* is modestly induced, mainly in the root epidermis, cortex, and root hair, and is detectable throughout the lateral root cap (Mudge *et al*., 2002, Karthikeyan *et al*., 2002). *AtPHT1;4* is highly induced by Pi limitation and expressed primarily in the epidermis, the cortex, and the root cap, but it is also present at the lateral root branch points on the primary root and in the stele of lateral roots (Misson *et al*., 2004). Besides the low Pi induction of *PHT1* genes at the transcriptional level, the trafficking of PHT1 proteins (PHT1s) to the PM is under constant surveillance. The ER exit of PHT1s requires PHOSPHATE TRANSPORTER TRAFFIC FACILITATOR1 (PHF1), an ER accessory protein sharing sequence homology with the small GTPase SAR1 guanine nucleotide exchange factor (GEF) SEC12 but missing the key catalytic residues required for the GEF activity (González *et al*., 2005). Dysfunction of *At*PHF1 specifically impairs Pi uptake by hampering the PM targeting of PHT1s, leading to a reduced abundance of PHT1s at the PM and a lower cellular Pi level, thereby activating Pi starvation-responsive (PSR) genes (González *et al*., 2005, Bayle *et al*., 2011). In rice, the phosphorylation status of the C-terminus of *Os*PHT1s affects their interaction with *Os*PHF1, and thus the ER exit of *Os*PHT1s (Chen *et al*., 2015, Yang *et al*., 2020). In addition, the turnover of PHT1s relies on the vacuolar degradation mediated either by the ENDOSOMAL SORTING COMPLEXES REQUIRED FOR TRANSPORT (ESCRT) III-related component (Cardona-López *et al*., 2015) or the ubiquitin E2 conjugase PHOSPHATE2 (PHO2) and/or the ubiquitin E3 ligase NITROGEN LIMITATION ADAPTATION (NLA) (Lin *et al*., 2013, Huang *et al*., 2013, Park *et al*., 2014). As a result, defective PHO2 or NLA confers Pi overaccumulation in plants due to increased accumulation of PHT1s (Lin *et al*., 2013, Huang *et al*., 2013).

Previous transcriptome profiling has uncovered *AtCNIH5* as a Pi starvation-induced gene with unknown implications (Müller *et al*., 2007, Liu *et al*., 2016). The membrane proteomics analysis of *Arabidopsis pho2* roots also revealed a significant increase in *At*CNIH5 (AT4G12090), along with a higher level of *At*PHT1;1–1;4 and *At*PHF1 (Huang *et al*., 2013). Although it remains to be determined whether *At*CNIH5 is a direct target of *At*PHO2, when querying a yeast split-ubiquitin-based *Arabidopsis* interactome database (*At*CNIH5 not included), we found that *At*CNIH1 interacts with *At*PHT1;9 and *At*PHF1 (Jones Alexander *et al*., 2014). These data prompted us to speculate that Pi starvation-induced *At*PHF1, *At*CNIH5, and *At*PHT1s are genetically and functionally linked. We were thus curious whether *At*CNIH proteins (*At*CNIHs), particularly *At*CNIH5, are involved in the regulatory mechanism for the post-ER trafficking of *At*PHT1s. In this study, we characterized *AtCNIH5* as the sole *AtCNIH* gene induced by Pi starvation in the outer root cell layers, including epidermal, cortical, and endodermal cells. We showed that in agro-infiltrated *Nicotiana benthamiana* (*N. benthamiana*) leaves, the fluorescent-tagged *At*CNIH5 is ER-localized and closely associated with the ER exit sites (ERES) markers *At*SAR1A, *At*SEC16A, and *At*SEC24A. We confirmed the interaction of *At*CNIH5 with *At*PHT1;1 using the *in-planta* split-GFP complementation and the yeast split-ubiquitin assays. The co-immunoprecipitation (co-IP) results also demonstrated that GFP-*At*CNIH5 interacts with the endogenous *At*PHT1;1/2/3 proteins. The *cnih5* mutant exhibits reduced shoot Pi levels, decreased *At*PHT1s, and increased *At*PHF1 accumulation. Consistently, the PM localization of split-GFP tagged*-At*PHT1;1 is diminished in the epidermis of the root transition/elongation zone and the root hair. The characterization of *cnih1/5*, *cnih3/5,* and *cnih4/5* double mutants further suggested that *At*CNIH5 is the primary *At*CNIH member regulating the cellular Pi homeostasis during the seedling stage. Moreover, our genetic studies suggest that dysfunctional *At*CNIH5 alleviates the Pi toxicity of the Pi overaccumulator *pho2*, supporting a role for *At*CNIH5 in regulating Pi homeostasis. Nevertheless, neither *At*CNIH5 interacts with *At*PHO2 *in planta*, nor does the transient expression of *At*PHO2 downregulate the expression of *At*CNIH5, arguing against *At*CNIH5 as a direct target of the *At*PHO2 regulatory module. Overall, our findings demonstrate that loss of *AtCNIH5* leads to a lower PM targeting efficiency of *At*PHT1s and reveal *At*CNIH5 as a *bona fide* ER cargo receptor that interplays with *At*PHF1 to ensure efficient trafficking of *At*PHT1s to the cell surface.

## Results

### *At*CNIH5 is induced by Pi deficiency preferentially in the outer root cell layers

Previous RNA-seq analysis (Liu *et al*., 2016) suggested that *AtCNIH* gene members exhibit comparable expression levels in the roots of young seedlings, except *AtCNIH2*, whose expression was barely detectable (Fig. S1). To corroborate the responsiveness of *AtCNIH5* to Pi starvation, we examined the transcript expression of *AtCNIH1, AtCNIH3, AtCNIH4*, and *AtCNIH5* by RT-qPCR. The expression of *AtCNIH5* remarkably increased in both the shoot and root following Pi starvation (**Fig. 1A**). In contrast, *AtCNIH3* and *AtCNIH4* were upregulated by nitrogen (N) deprivation (**Fig. 1A**). We analysed the spatial expression pattern of *AtCNIH5* using promoter: *uidA* (a gene encoding β-glucuronidase or GUS) fusion lines. Under full-nutrient conditions, the promoter activity of *AtCNIH5* was found in the trichomes and the vascular tissues of the shoot (**Fig. 1B**). In the primary root, the expression of *AtCNIH5* was absent in the root cap and meristem but strongly expressed in the stele of the transition/elongation and nearly all cell types in the differentiation/maturation zones (**Fig. 1B**). In the lateral root, *AtCNIH5* was expressed in the vascular tissues and detectable in the columella (**Fig. 1B**). Interestingly, the promoter activity of *AtCNIH5* was markedly elevated in the epidermal, cortical, and endodermal cells of the primary root following Pi deprivation (**Fig. 1B**). In comparison, the promoter activities of *AtCNIH1*, *AtCNIH3*, and *AtCNIH4* were confined to the vascular tissues of the primary root, regardless of nutrient conditions (Fig. S2A), and also observed in the columella of the lateral root (Fig. S2B). Characterization of *AtCNIH5pro*: GFP fusion lines at the young seedling stage recapitulated that, irrespective of Pi regimes, *AtCNIH5* is expressed in the root transition/elongation and differentiation/maturation zones and root hairs (Fig. S3). Collectively, the expression of *AtCNIH5* is enhanced by Pi starvation, and its promoter activity is upregulated by low Pi in the outer root cell layers.

**Fig. 1.**
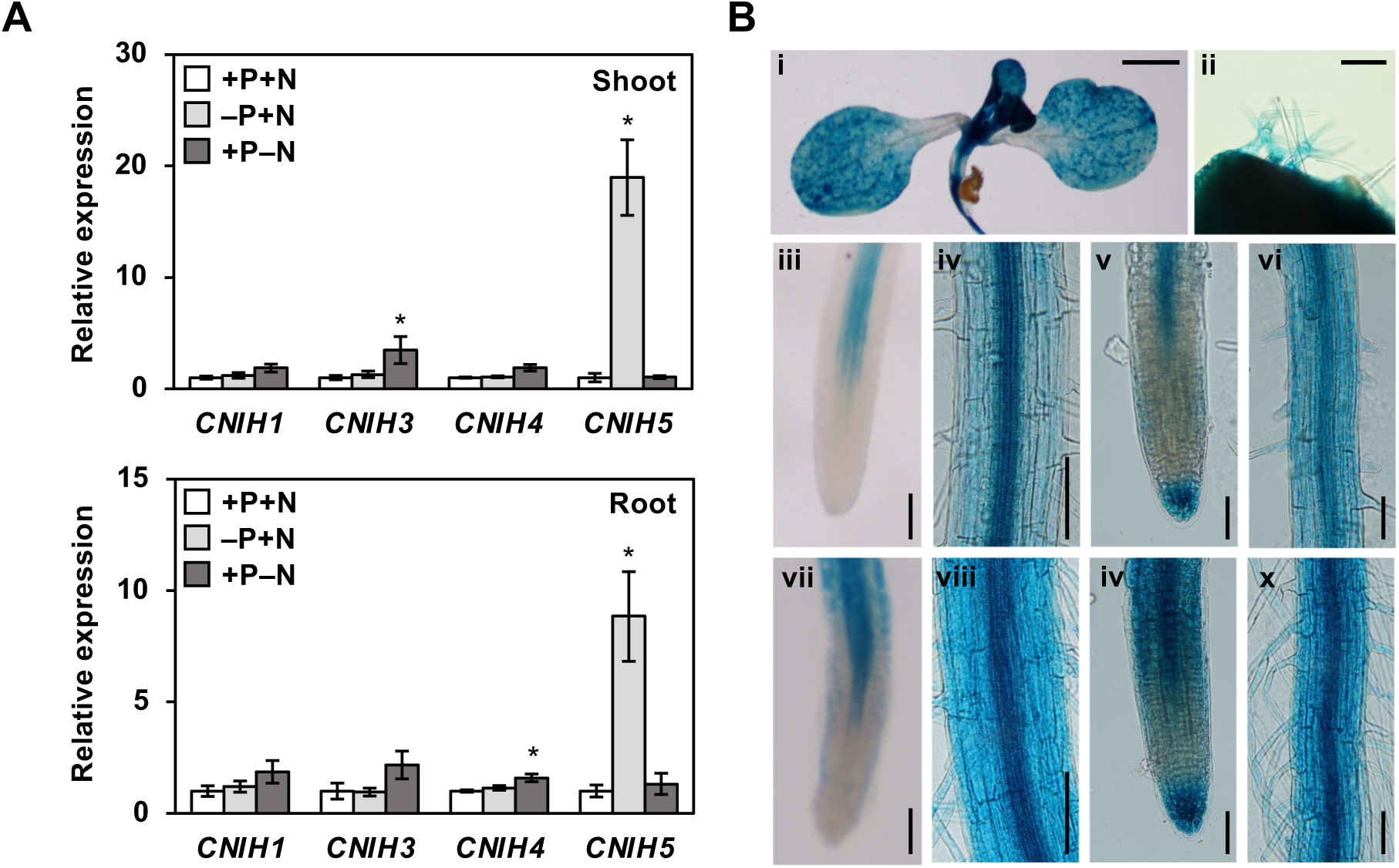
Pi starvation-induced expression pattern of *At*CNIH5. (A) RT-qPCR analysis of *A. thaliana CNIH1, CNIH3, CNIH4,* and *CNIH5* in the shoot and root of 11-day-old seedlings grown under +P+N (250 µM KH_2_PO_4_ and 7.5 mM KNO_3_), –P+N (0 µM KH_2_PO_4_ and 7.5 mM KNO_3_, three days of starvation), or +P–N (250 µM KH_2_PO_4_ and 0 µM KNO_3_, three days of starvation) conditions. Relative gene expression is presented as fold change compared to the control under +P+N conditions. Error bars represent SE (n = 4, pools of seedings from independent experiments). Dunn’s test following Kruskal–Wallis test, \**p* < 0.05. (B) Expression of *AtCNIH5pro*: GUS in 7-day-old seedlings grown under +P (250 µM KH_2_PO_4_) or –P (0 µM KH_2_PO_4_, five days of starvation) conditions. i–vi (+P; shoot part; trichome; primary root tip; primary root maturation zone; lateral root tip; lateral root maturation zone). vii–x (–P; primary root tip; primary root maturation zone; lateral root tip; lateral root maturation zone). Scale bars, 10 mm in i, 0.1 mm in ii, and 100 µm in iii–x. Representative images are shown from three independent T3 homozygous lines.

### *At*CNIH5 resides both at the ER and ERES

Previous studies of the pollen-specific expression of *At*CNIHs showed their distribution at the ER (Wudick *et al*., 2018). Furthermore, RFP-*At*CNIH4 could be detected at the pollen tube’s PM and colocalized with the ERES marker *At*SEC24A (Wudick *et al*., 2018). To examine the subcellular distribution of *At*CNIH5 in other tissues, we generated the construct encoding UBQ10 promoter-driven β-estradiol-inducible *At*CNIH5 tagged with the 10th β-strand of GFP (*UBQ10*:*sXVE*: S10-*At*CNIH5) for transient expression. In *N. benthamiana* leaves, the split-GFP self-complementation allows live-cell imaging of S10-tagged fusion proteins in the presence of the detector S11-GFP1–9 (Liu *et al*., 2018, Liu, 2021). The distribution of S10-*At*CNIH5 showed a typical ER-network pattern when co-expressed with the cytosolic S11-GFP1–9 (**Fig. 2A**), indicating that the N-terminus of *At*CNIH5 is in the cytoplasm. While S10-*At*CNIH5 colocalized with the ER-luminal marker mCherry-HDEL, the mCherry-*At*CNIH5-labeled punctate structures distributed on the ER tubules and sheet rims and were closely associated with the ERES marker *At*SEC16A/MAG5 (Takagi *et al*., 2013) (**Fig. 2A**). Taking advantage of high-resolution confocal microscopy, we measured the distance between the red fluorescence protein-tagged *At*CNIH5 and the *cis*-Golgi marker MNS1 (Liebminger *et al*., 2009) and the three ERES markers *At*SAR1A-GFP, GFP-*At*SEC16A, and GFP-*At*SEC24A. Our results demonstrated that both the constitutively expressed RFP-*At*CNIH5 and the inducibly expressed mCherry-*At*CNIH5 significantly colocalized with the tested ERES markers but not with the *cis*-Golgi marker MNS1-eGFP (**Fig. 2B**, **2C**).

**Fig. 2.**
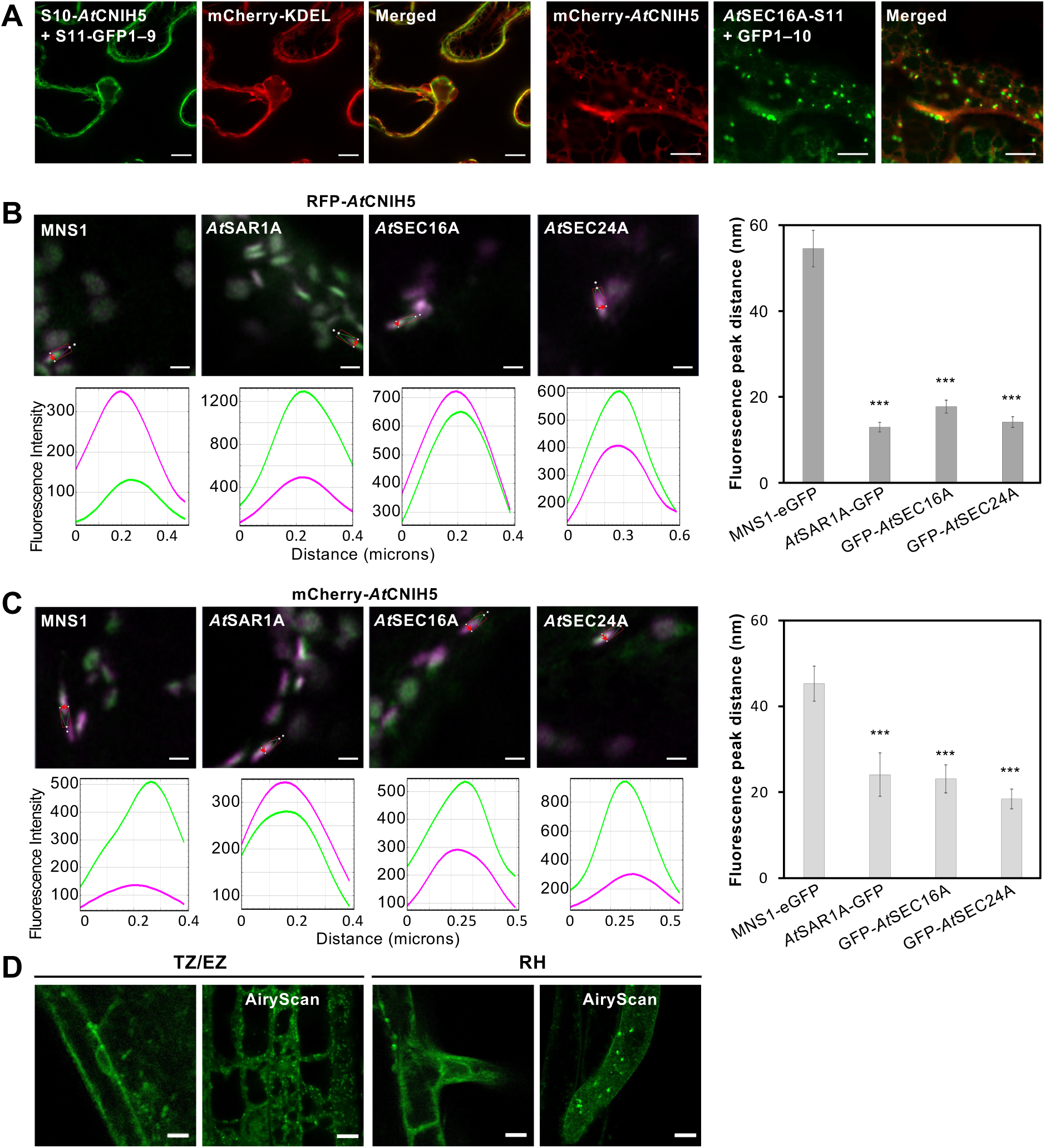
Subcellular localization of *At*CNIH5 to the ER and ERES. (A) The ER and ERES localizations of S10-*At*CNIH5 in agro-infiltrated *N. benthamiana* leaves. mCherry-HDEL, luminal ER marker; *At*SEC16A-S11, ERES marker. Images are from single confocal slices. Scale bars, 10 µm. (B, C) Statistical line profile analysis of fluorescence intensity for localization of *At*CNIH5 with the ERES and *cis*-Golgi markers. Representative images and line profile analysis for *35S:* RFP-*At*CNIH5 (B) and *UBQ10*:*sXVE*: mCherry-*At*CNIH5 (C) (magenta) with the *cis*-Golgi marker MNS1-eGFP and the ERES markers *At*SAR1A-GFP, GFP-*At*SEC16A, and GFP-*At*SEC24A (green). The line profile applied and the line profile output are shown. Peak distance analysis graphs showing the co-localization analysis with RFP-*At*CNIH5 and mCherry-*At*CNIH5 with the ERES and *cis*-Golgi markers. Scale bars = 1 µm. Kruskal–Wallis analysis was used to compare the median peak distances among multiple groups. Post-hoc pairwise comparisons were performed using Dunn’s multiple comparison test with Bonferroni correction to identify significant group differences (\*\*\**p* < 0.001). Error bars represent SE (n = 4 biological replicas, with at least 18 technical repeats). (D) Localization of GFP-*AtCNIH5* in the root of *Arabidopsis* 5-day-old seedlings grown under –P (0 µM KH_2_PO_4_, five days of starvation) conditions. TZ/EZ, transition zone/elongation zone; RH, root hair. Images are from single confocal slices. Scale bars, 10 µm.

To validate the subcellular localization of *At*CNIH5 in stable lines, we used *cnih5* as the genetic background to obtain transgenic plants carrying the genomic sequence of *AtCNIH5* translationally in-frame with GFP (*AtCNIH5pro*: GFP-*AtCNIH5*). Two independent lines showed a modest increase in shoot biomass and restored the shoot Pi content to wild-type (WT)-like levels, while their shoot Pi concentrations were lower—though not significantly—than those of the *cnih5* mutant (Fig. S4A), suggesting an inverse relationship between shoot fresh weight and Pi levels in these lines. After enriching microsomal fractions, we also detected the expression of GFP-*AtCNIH5* proteins in Pi-limited roots using anti-*At*CNIH5 antibody (Fig. S4C). However, it is technically challenging to determine whether GFP-*AtCNIH5* proteins are more abundant than the endogenous *At*CNIH5 proteins in WT plants due to differences in molecular weight between GFP-*AtCNIH5* and *At*CNIH5. Under the confocal microscope, the signals of GFP-*AtCNIH5* in the Pi-limited roots appeared faint overall. Nonetheless, when images were acquired using Airyscan mode, the punctate structures in the root hair and root epidermal cells became more pronounced (**Fig. 2D**). These observations suggested that the localization of *At*CNIH5 at the ER and ERES in *N. benthamiana* leaves was unlikely to be an artifact of overexpression.

### *At*CNIH5 interacts with *At*PHT1s and *At*PHF1

The yeast split ubiquitin-based interactome revealed the interaction of *At*CNIH1 with *At*PHT1;9 and *At*PHF1 (Jones Alexander *et al*., 2014), prompting us to test whether *At*CNIH5 also interacts with *At*PHT1s and *At*PHF1. Considering the expression of *AtCNIH5* and *AtPHT1;1* mainly in the root epidermis, we selected *At*PHT1;1 to represent *At*PHT1s in this study. In agro-infiltrated *N. benthamiana* leaves, S10-*At*CNIH5 interacted with the 11th β-strand of sfGFP-tagged *At*PHT1;1 (*At*PHT1;1-S11), as manifested by the reconstituted GFP signals in the presence of the cytosolic GFP1–9 (Liu *et al*., 2018, Liu, 2021) (**Fig. 3A**). *At*CNIH5 and *At*PHF1 have a similar membrane topology, making the tripartite split-GFP complementation method unsuitable for detecting their interaction. Therefore, we used the conventional bipartite split-GFP complementation method, known as bimolecular fluorescence complementation (BiFC), to investigate their interaction. However, we were unable to demonstrate an interaction between these two proteins because positive controls failed. We then exploited the yeast split-ubiquitin system (SUS) to examine the interaction of *At*CNIH5 with *At*PHT1;1, *At*PHT1;4, and *At*PHF1. The results of the nutrient selection and the LacZ β-galactosidase assays suggested that *At*CNIH5 interacted with these proteins (**Fig. 3B**). Since the amino acid sequences of *At*PHT1s share over 47.4% identity, we demonstrated that *At*CNIH5 also interacts with *At*PHT1;2, *At*PHT1;5, *At*PHT1;7 and *At*PHT1;9 in our side-by-side study (Liu *et al*., 2024), indicating that *At*CNIH5 recognizes conserved motifs within the *At*PHT1 family. Furthermore, we performed co-IP using *Arabidopsis* transgenic plants harbouring the genomic GFP-*AtCNIH5* (*AtCNIH5pro*: GFP-*AtCNIH5/cnih5*) or the free GFP under the UBQ10 promoter. As the cytosolic GFP was nearly 20 times more abundant than GFP-*AtCNIH5* in the root low-speed pellet (LSP) membrane fraction, we calculated that approximately 40 times more endogenous *At*PHT1;1/2/3 and *At*PHF1 proteins were co-immunoprecipitated with GFP-*AtCNIH5* than with GFP (**Fig. 3C**). These results supported that *At*CNIH5 interacts with *At*PHT1s and *At*PHF1.

**Fig. 3.**
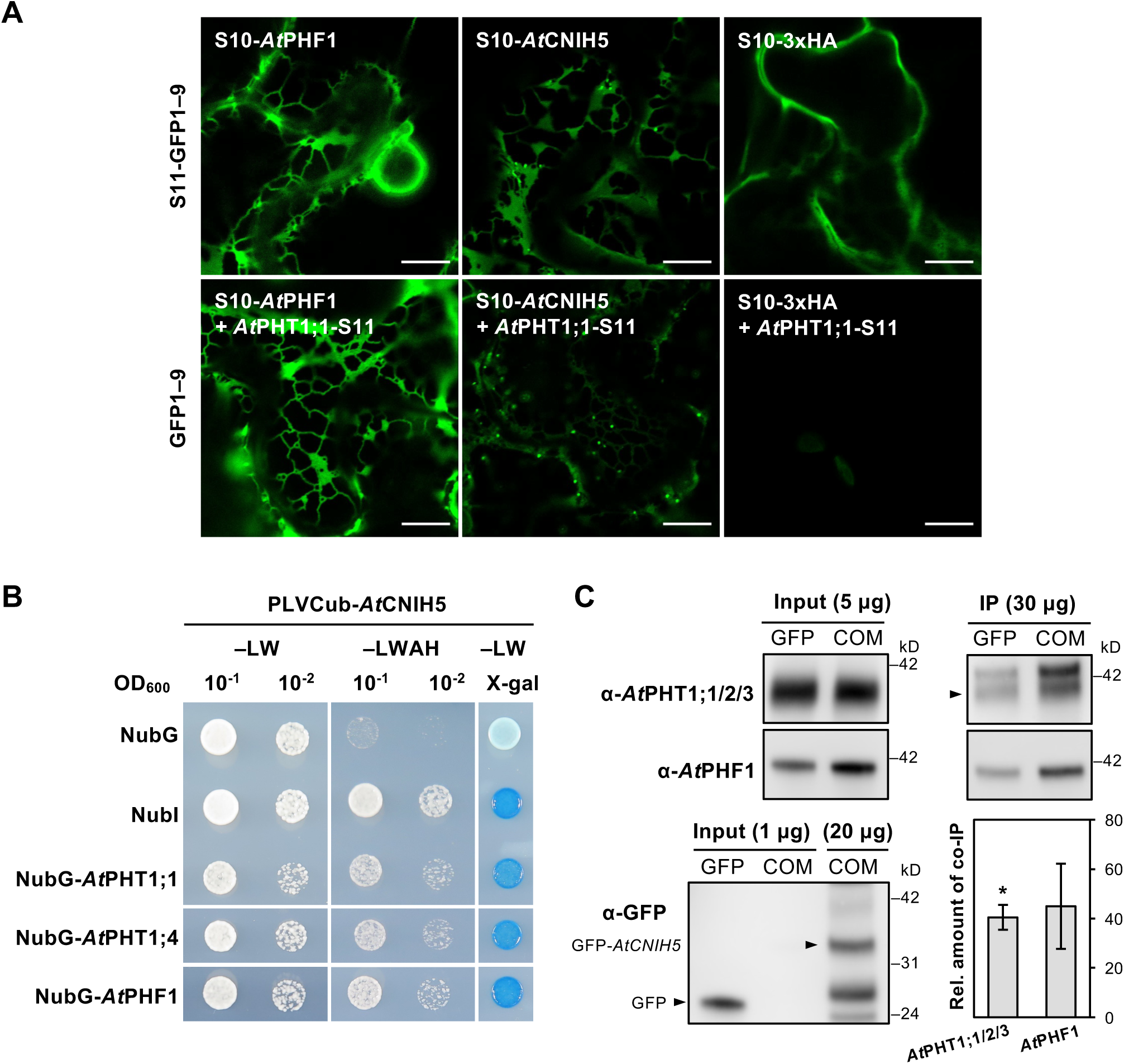
*At*CNIH5 interacts with *At*PHT1;1, *At*PHT1;4, and *At*PHF1. (A) Expression of S10-*At*CNIH5, S10-*At*PHF1, S10-3xHA and *At*PHT1;1-S11 in the presence of S11-GFP1–9 or GFP1–9 in *N. benthamiana* leaves. The co-expression of *At*PHT1;1-S11 with S10-*At*PHF1 and S10-3xHA are used as positive and negative protein–protein interaction controls, respectively. Images are from single confocal slices. Scale bars, 10 µm. (B) Co-expression of PLVCub-*At*CNIH5 with NubG-*At*PHF1, NubG-*At*PHT1;1, and NubG-*At*PHT1;4 in the yeast split-ubiquitin system. The co-expression of PLVCub-*At*CNIH5 with NubI and NubG are used as positive and negative controls, respectively. Yeast transformants were grown on synthetic medium without leucine and tryptophan (–LW; the left panel) or on synthetic medium lacking leucine, tryptophan, adenine, and histidine-containing 500 μM methionine (–LWAH; middle panel) or on SC–LW containing 2 mg/L X-gal (X-gal; the right panel). (C) Coimmunoprecipitation of *At*PHT1;1/2/3 and *At*PHF1 with GFP-*AtCNIH5* in 11-day-old *AtCNIH5pro*: GFP-*AtCNIH5/cnih5* complementation (COM) seedlings grown under –P (0 µM KH_2_PO_4_, seven days of starvation) conditions. A *UBQ10*: GFP line was used as a control. The amount of root low-speed pellet (LSP) membrane fraction used for the input or the immunoprecipitation (IP) loading is shown as indicated. The relative abundance of co-immunoprecipitated endogenous *At*PHT1;1/2/3 and *At*PHF1 proteins was calculated by normalizing them with the GFP control. Data significantly different from the GFP control are indicated by asterisks (n = 3 biological replicates; \**p* < 0.05; Student’s t-test, two-tailed).

### Loss of *AtCNIH5* confers reduced Pi levels, *At*PHT1s accumulation, and Pi transport

To determine the physiological role of *At*CNIH5, we grew WT and *cnih5* mutants under different Pi regimes. Regardless of Pi status, the shoot and root fresh weight of *cnih5* mutants showed no differences from that of WT (**Fig. 4A**). In contrast, the shoot Pi level of *cnih5* diminished by 18% compared to WT under Pi sufficiency (**Fig. 4B**). We could rescue both the lower shoot Pi concentrations and Pi content in *cnih5* under Pi sufficiency by introducing the genomic construct encoding *At*CNIH5 (*AtCNIH5pro*: *AtCNIH5*) (Fig. S4B), indicating that the reduced Pi accumulation in *cnih5* was attributed to the non-functional *cnih5* allele. While the protein expression of the *AtCNIH5* transgene in the two independent lines was similar to the endogenous *At*CNIH5 levels in WT, the third line exhibited a 4.4-fold increase in *At*CNIH5 protein levels and grew slightly larger than WT (Fig. S4B, S4C). These findings are consistent with the restored shoot Pi content *per* plant and the inverse relationship between the shoot fresh weight and the shoot Pi levels noted in the GFP-*AtCNIH5*-complemented lines (Fig. S4A). We inferred that the increased shoot biomass of GFP-*AtCNIH5* complemented lines was likely due to the enhanced expression or activity of GFP-*AtCNIH5* compared to that of the genomic *AtCNIH5* transgene.

**Fig. 4.**
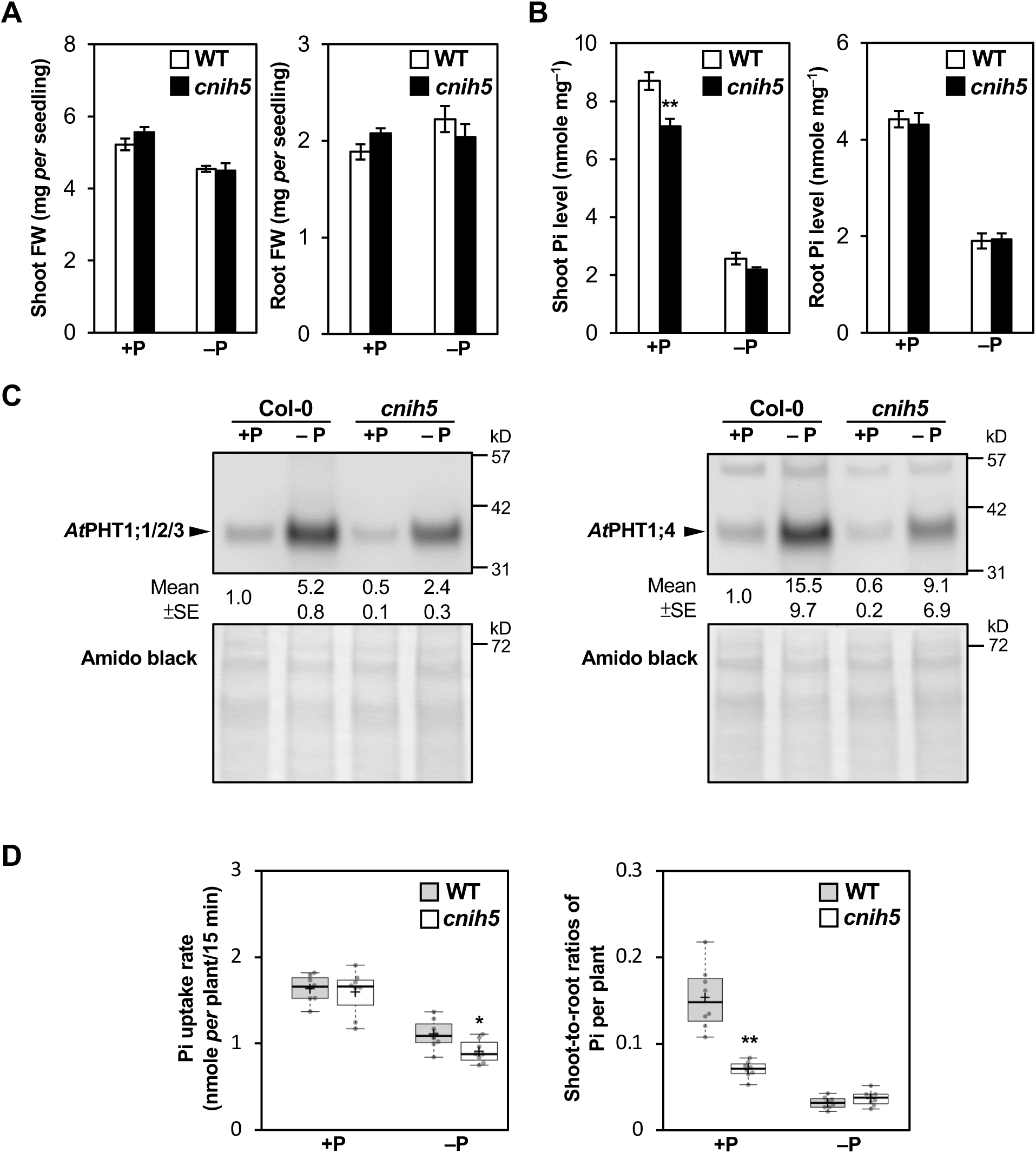
Loss of *AtCNIH5* leads to decreased shoot Pi levels, lower abundance of *At*PHT1 proteins, and diminished Pi uptake. (A, B) The shoot and root fresh weight (FW) (A) and Pi levels (B) of 11-day-old *Arabidopsis* WT and *cnih5* seedlings under +P (250 µM KH_2_PO_4_) or–P (0 µM KH_2_PO_4_, three days of starvation) conditions. Error bars represent SE (n = 15–16, pools of seedlings from independent experiments; \**p* < 0.05, \*\**p* < 0.01; Student’s t-test, two-tailed). (C) Expression of *At*PHT1;1/2/3 and *At*PHT1;4 in the root microsomal fractions from 11-day-old *Arabidopsis* WT and *cnih5* seedlings under +P (250 µM KH_2_PO_4_) or –P (0 µM KH_2_PO_4_, three days of starvation) conditions. The relative expression levels of *At*PHT1;1/2/3 and *At*PHT1;4 were normalized with the corresponding amido black staining and relative to the WT control. Error bars represent SE (n = 2, pools of seedlings from independent experiments). (D) Quantification of Pi uptake and shoot-to-root distribution. Data are visualized using BoxPlotR (Spitzer *et al*., 2014). The centre lines show the medians; the central plus signs (+) show the means; box limits indicate the 25th and 75th percentiles; whiskers extend to the minimum and the maximum values. Data points are plotted as dots. Data significantly different from WT are indicated by asterisks (n = 8 biological replicates; \**p* < 0.05, \*\**p* < 0.01; Student’s t-test, two-tailed).

The lower shoot Pi levels in *cnih5* did not lead to the upregulation of several Pi starvation-induced genes in the root, except for *AtPHT1;4* (Fig. S5A). The primary root length was slightly shorter in *cnih5* than in WT regardless of Pi regimes, while the lateral root density in *cnih5* was significantly higher than in WT under Pi sufficiency (Fig. S5B, S5C). In addition, *cnih5* exhibited shorter root hair lengths than WT, but there was no difference in root hair density between WT and *cnih5* (Fig. S5E, S5F). To evaluate whether *At*PHT1s account for the decreased Pi levels in *cnih5*, we examined the abundance of *At*PHT1;1/2/3 and *At*PHT1;4 proteins, which are the prominent *At*PHT1 members involved in Pi uptake (Shin *et al*., 2004, Ayadi *et al*., 2015). Given that the impaired ER exit of *At*PHT1s in *phf1* caused a reduction in *At*PHT1;1/2/3 and *At*PHT1;4 proteins (Huang *et al*., 2013), we hypothesized that PM proteins retained inside the cell are eventually targeted for degradation due to protein quality control. Based on this, defective *At*CNIH5 would reduce the steady-state levels of *At*PHT1;1/2/3 and *At*PHT1;4. Although the expression of *AtCNIH1*, *AtCNIH3*, and *AtCNIH4* was of a similar magnitude in our qRT-PCR results (**Fig. 1A**), previous RNA-seq analysis (Liu *et al*., 2016) suggested *AtCNIH4* as the most abundant *AtCNIH* gene member (Fig. S1). Therefore, we included *cnih4* mutants for comparison. Under Pi sufficiency, the amount of *At*PHT1;1/2/3 and *At*PHT1;4 in the total root proteins was indistinguishable among WT, *cnih4*, and *cnih5*; however, under Pi deficiency, these proteins were decreased in *cnih5* by 20–25% as compared to WT (Fig. S6). After enrichment of root membrane proteins, a 50% decrease of *At*PHT1;1/2/3 was detected in *cnih5* under both Pi regimes (**Fig. 4C**). The protein abundance of *At*PHT1;4 was also reduced by 40% in the root membrane proteins of *cnih5* as compared with the corresponding WT controls (**Fig. 4C**). Importantly, the protein abundance of *At*PHT1;1/2/3 in the Pi-limited *cnih5* complemented lines either carrying *AtCNIH5pro*: GFP-*AtCNIH5* or *AtCNIH5pro*: *AtCNIH5* were higher than in *cnih5* and were restored to levels comparable to or exceeding those of the WT (Fig. S4C), reinforcing that *At*PHT1;1/2/3 are indeed downstream effectors of *At*CNIH5.

We determined the Pi uptake of radioactive tracer (^32^P) *per* plant using a real-time imaging system (Kanno *et al*., 2016). When Pi is replete, there were no significant differences in the Pi uptake within 15 minutes between WT and *cnih5*; however, the shoot-to-root distribution of ^32^P significantly decreased in *cnih5* (**Fig. 4D**). This may be attributed to an additional role of *At*CNIH5 in the translocation of Pi from roots to shoots under Pi sufficiency by facilitating the PM targeting of *At*PHT1 transporters in different root cell types, as several *AtPHT1* genes are expressed in the vascular tissues (Nussaume *et al*., 2011). Under Pi deficiency, loss of *AtCNIH5* resulted in an 18% reduction of ^32^P uptake relative to the WT, while the shoot-to-root distribution of ^32^P was comparable between the two genotypes (**Fig. 4D**), suggesting that *At*CNIH5 is more critical for Pi uptake under Pi limitation. Overall, *cnih5* exhibited lower shoot Pi accumulation under Pi sufficiency due to the reduced translocation of Pi from roots to shoots and showed decreased Pi uptake under Pi deficiency, which could be ascribed to the decreased protein amount of *At*PHT1;1/2/3 and *At*PHT1;4, and probably other *At*PHT1s as well.

To answer whether *At*CNIH5 shares functional redundancy with other *At*CNIH members, we analysed *cnih* double mutants carrying defective *AtCNIH1*, *AtCNIH3*, or *AtCNIH4* in the *cnih5* background. In the respective *cnih* double mutants, the expression of the remaining *AtCNIH* genes was not altered (Fig. S7A), hinting that there is no compensatory upregulation of other *AtCNIH* genes at the transcript level when *AtCNIH5* and/or other *AtCNIH* are knocked out. Compared to *cnih5*, all the *cnih* double mutants did not display a greater reduction in fresh weight and Pi levels (Fig. S7B, S7C). Taken together, *At*CNIH5 is the predominant *At*CNIH member functionally linked to the regulation of Pi transport.

### Loss of *AtCNIH5* leads to decreased plasma membrane localization of *At*PHT1;1

To prove that *At*CNIH5 is a *bona fide* ER cargo receptor of *At*PHT1s, we sought to check the subcellular distribution of *At*PHT1;1 in *cnih5*. For equal transgene expression, a homozygous line (*At*PHT1;1-S11/GFP1–10) co-expressing *At*PHT1;1-S11 driven by the native *At*PHT1;1 promoter and GFP1–10 driven by the 35S promoter was generated in the WT background and outcrossed to the *cnih5* mutant. The F2 progeny were screened for moderate fluorescence signals and subjected to genotyping. F3 individuals propagated from the F2 plants with either the WT (+/+) or *cnih5* (–*/*–) background were used for confocal imaging analysis. Under Pi sufficiency, the fluorescence signals of *At*PHT1;1-S11/GFP1–10 plants in the WT background were relatively weak. Therefore, we focused on the subcellular localization of *At*PHT1;1-S11 in the area where *AtPHT1;1* is expressed under Pi limitation. In the WT background, *At*PHT1;1-S11 accumulated at the PM in the root epidermis of the transition/elongation zone (**Fig. 5A**, **5B**) and the root hairs of the differentiation/maturation zone (**Fig. 5C**). By contrast, *At*PHT1;1-S11 in the *cnih5* background occasionally distributed into the intracellular reticular or punctate structures in these regions (**Fig. 5B**, **5C**, and **Table 1**), and the fluorescence signals significantly declined at the PM by nearly 50% (**Fig. 5A**, **5C**). Hence, we conclude that *At*CNIH5 facilitates the ER export and thus the PM targeting of *At*PHT1;1, particularly in the elongating root epidermis and root hair.

**Fig. 5.**
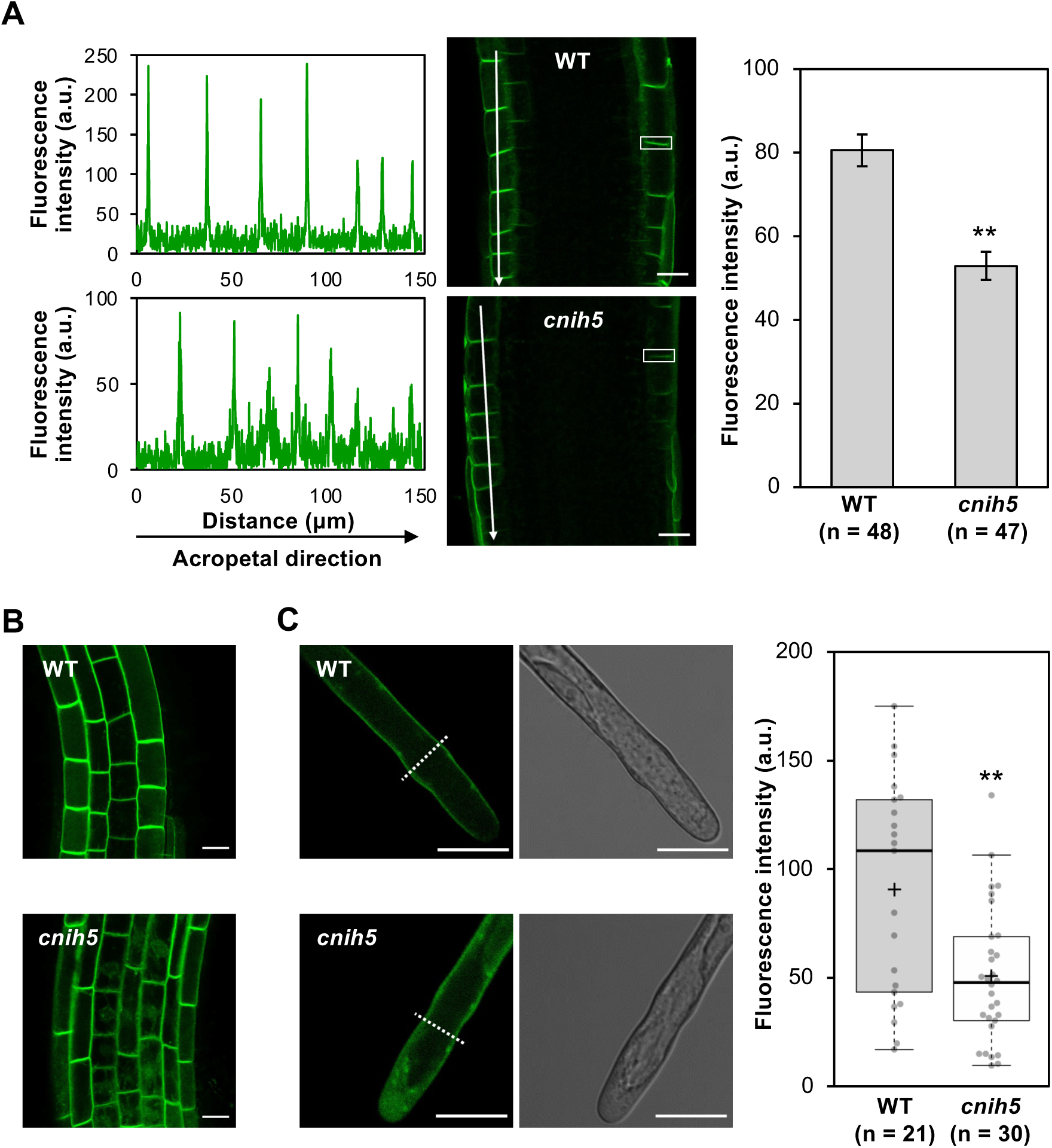
Impaired plasma membrane targeting of split-GFP-tagged *At*PHT1;1 in *cnih5* root. (A) Reduction of *At*PHT1;1-S11 at the plasma membrane of the epidermal cells in the root transition/elongation zone of 5-day-old *cnih5* seedlings under −P (0 μM KH_2_PO_4_) conditions. The transgenic lines expressing *AtPHT1;1pro*: *At*PHT1;1-S11/*35S*: GFP1–10 in the *cnih5* and corresponding WT backgrounds were used for comparison. Representative images and line-scan fluorescence intensity profiles (arbitrary unit, a.u.) of *At*PHT1;1-S11 are shown in the left panel. Arrows indicate the acropetal direction. The fluorescence intensity histogram of *At*PHT1;1-S11 enclosed by the representative rectangle is quantified in the right panel. Data significantly different from WT are indicated by asterisks (n, counted regions from transition/elongation zone of 8 roots; \*\**p* < 0.01; Student’s t-test, two-tailed). Scale bars, 20 µm. (B) A representative image shows increased intracellular retention of *At*PHT1;1-S11 in the root epidermal cells along the transition zone and the elongation zone of *cnih5*. Scale bars, 20 µm. (C) Reduction of *At*PHT1;1-S11 at the plasma membrane in the *cnih5* root hair. Line-scan profiles were performed at a distance of 30 µm from the root hair tip as indicated by the dashed line for quantification of PM fluorescence intensity (arbitrary unit, a.u.). All the fluorescence images are acquired from a single confocal optical slice. Quantitative data in the right panel are visualized using BoxPlotR (Spitzer *et al*., 2014). The centre lines show the medians; the central plus signs (+) show the means; box limits indicate the 25th and 75th percentiles; whiskers extend to the smallest data point within 1.5 times the interquartile range from the lower and upper quartiles. Data significantly different from WT are indicated by asterisks (n, counted root hairs; \*\**p* < 0.01; Student’s t-test, two-tailed). Scale bars, 20 µm.

**Table 1.** Intracellular retention of split-GFP tagged*-At*PHT1;1 in the root transition/elongation zones of *cnih5*.

| | | | Root 1 | Root 2 | Root 3 | Root 4 | Average (%) ± SE<br>(n=4) | *P*-value |
| --- | --- | --- | --- | --- | --- | --- | --- | --- |
|  |  |  | No. of cells with intracellular PHT1;1 |  |  |  |  |  |
|  |  |  | /No. of counted cells |  |  |  |  |  |
| TZ/EZ | Exp. 1 | WT | 0/9 | 0/13 | 5/18 | 1/12 | 9.0±6.6 | <0.001 |
| | | *cnih5* | 10/17 | 9/15 | 11/15 | 8/12 | 64.7±3.4 | |
| | Exp. 2 | WT | 1/12 | 1/13 | 0/7 | 0/9 | 4.0±2.3 | <0.001 |
| | | *cnih5* | 7/12 | 4/8 | 7/9 | 5/6 | 67.4±7.9 | |

- TZ, transition zone; EZ, elongation zone. - Exp. 1 and Exp. 2 represent two independent experiments. In each experiment, four seedlings expressing *AtPHT1;1*-S11 in either the WT or *cnih5* background were used for image analysis. - Average (%) ± SE (n = 4) indicates the mean percentage of cells containing intracellular *AtPHT1;1*-S11 among those counted, along with the standard error of the mean. - *P*-value is calculated based on a two-tailed Student's t-test.

### *At*CNIH5 and *At*PHF1 act in concert to control the ER exit of *At*PHT1s

The ER-resident *At*PHF1 was previously known to regulate the ER exit of *At*PHT1s. We thus conjectured whether the decreased PM targeting of *At*PHT1s in *cnih5* is indirectly due to an altered protein abundance of *At*PHF1. To our surprise, *At*PHF1 increased by nearly 1.3-fold in the root total proteins of *cnih5* under both Pi regimes (Fig. S6), even though the transcript level of *AtPHF1* was not upregulated (Fig. S5A). These results implied that *At*PHF1 is likely upregulated to compensate for the loss of *At*CNIH5. Considering that *At*CNIH5, as well as *At*PHF1, interacts with *At*PHT1;1, we suspected that *At*CNIH5 collaborates with *At*PHF1 to assist the ER exit of *At*PHT1s. To explore this possibility, we characterized *phf1/cnih4*, *phf1/cnih5*, and *phf1/cnih4/5* mutants. Compared to the *phf1* seedlings, the shoot and root biomass of *phf1/cnih5* and *phf1/cnih4/5* mutants were further reduced (**Fig. 6A**). However, *phf1/cnih5* mutants showed no differences in the shoot and root Pi levels compared to the *phf1* (**Fig. 6B**), suggesting that *phf1* is epistatic to *cnih5*. In accordance with these results, the protein amount of *At*PHT1;1/2/3 in the total root proteins was comparable in these mutants (**Fig. 6C**). During the adult-plant stage in hydroponic solutions, the *phf1/cnih4*, *phf1/cnih5*, and *phf1/cnih4/5* mutants also displayed reduced shoot growth relative to the *phf1* (Fig. S8A). Of note, the *phf1/cnih5* and *phf1/cnih4/5* plants exhibited reduced shoot Pi levels relative to the *phf1* under Pi sufficiency (Fig. S8B). Therefore, the extra loss of *AtCNIH5* in the *phf1* background aggravates the growth retardation and further reduces shoot Pi accumulation when plants grow to an older stage, indicating a complex genetic interaction between *At*PHF1 and *At*CNIH5.

**Fig. 6.**
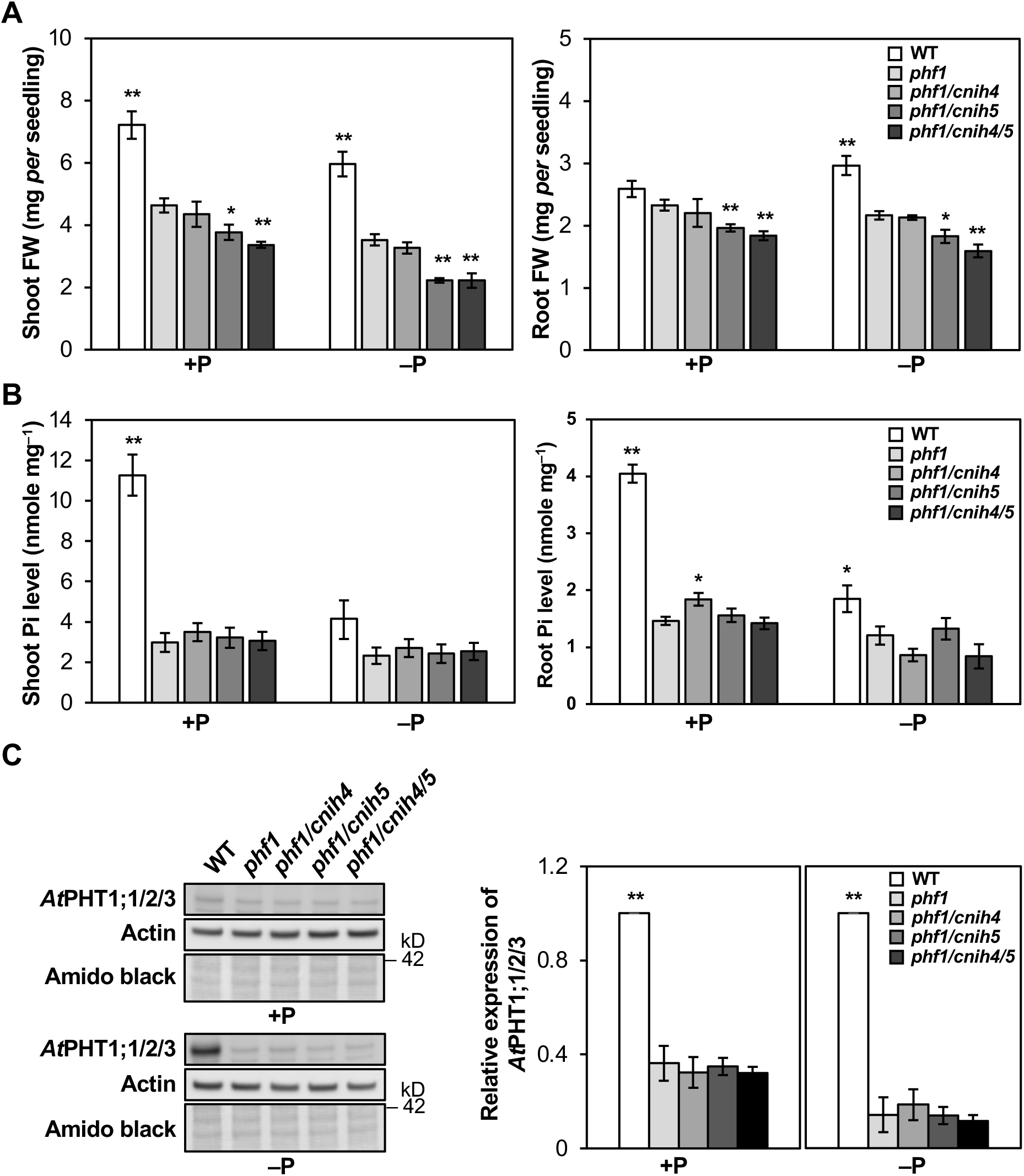
Loss of *AtCNIH5* exacerbates the growth retardation of *phf1*. (A, B) The shoot and root fresh weight (FW) (A) and Pi levels (B) of 11-day-old *Arabidopsis* WT, *phf1, phf1/cnih4, phf1/cnih5*, and *phf1/cnih4/5* seedlings grown under +P (250 µM KH_2_PO_4_) or –P (0 µM KH_2_PO_4_, three days of starvation) conditions. Error bars represent SE (n = 7–8, pools of seedlings from independent experiments. (C) The expression of *At*PHT1;1/2/3 in the root of 11-day-old *Arabidopsis* WT, *phf1, phf1/cnih4, phf1/cnih5*, and *phf1/cnih4/5* seedlings grown under +P (250 µM KH_2_PO_4_) or –P (0 µM KH_2_PO_4_, three days of starvation) conditions. The relative expression level of *At*PHT1;1/2/3 was normalized with the corresponding actin and relative to the WT control. Error bars represent SE (n = 3, pools of seedlings from independent experiments). One-way ANOVA with post-hoc Dunnett’s test versus *phf1,* \**p* < 0.05, \*\**p* < 0.01.

### *At*CNIH5 alleviates the Pi toxicity of *pho2* but is unlikely to be a direct target of *At*PHO2

To address whether *At*CNIH5 acts downstream of the *At*PHO2-mediated Pi homeostasis pathway, we generated the *pho2/cnih5* mutant. Compared to *pho2*, *pho2/cnih5* accumulated less Pi in the shoot to a similar extent as reported for *pho2/pht1;1* (**Fig. 7A**) (Huang *et al*., 2013) and exhibited reduced levels of *At*PHT1;1/2/3 and *At*PHT1;4 relative to the *pho2* (**Fig. 7B**). By comparison, the protein level of the xylem loading Pi transporter *At*PHO1 was increased in *pho2/cnih5* (**Fig. 7B**). The upregulation of *At*PHO1 was also observed in the *cnih5* mutant regardless of Pi regimes (Liu *et al*., 2024), suggesting that defective *AtCNIH5* specifically impairs the ER export of *At*PHT1s. We next explored whether *At*PHO2 functions as a hypothesized E2/E3 hybrid enzyme to mediate the downregulation of *At*CNIH5. It has been shown that the C748A mutation of *At*PHO2 disrupts the conserved catalytic cysteine residue necessary for *At*PHO2-mediated degradation of *At*PHO1 (Liu *et al*., 2012). Therefore, we utilized this variant in the protein-protein interaction assay to avoid substrate or protein degradation. Using the *N. benthamiana* transient expression system, we found that, in contrast to *At*PHO2-mediated degradation of *At*PHO1 in a dosage-dependent manner as previously reported (Liu et al., 2012), neither the catalytically defective *At*PHO2^C748A^ interacted with *At*CNIH5 nor the catalytically active *At*PHO2 suppressed the expression of *At*CNIH5 (Fig. S9A, S9B). Taken together, the loss of *AtCNIH5* alleviates the Pi toxicity of *pho2* due to downregulation of *At*PHT1s, but we have no evidence to support *At*CNIH5 as a direct target of *At*PHO2.

**Fig. 7.**
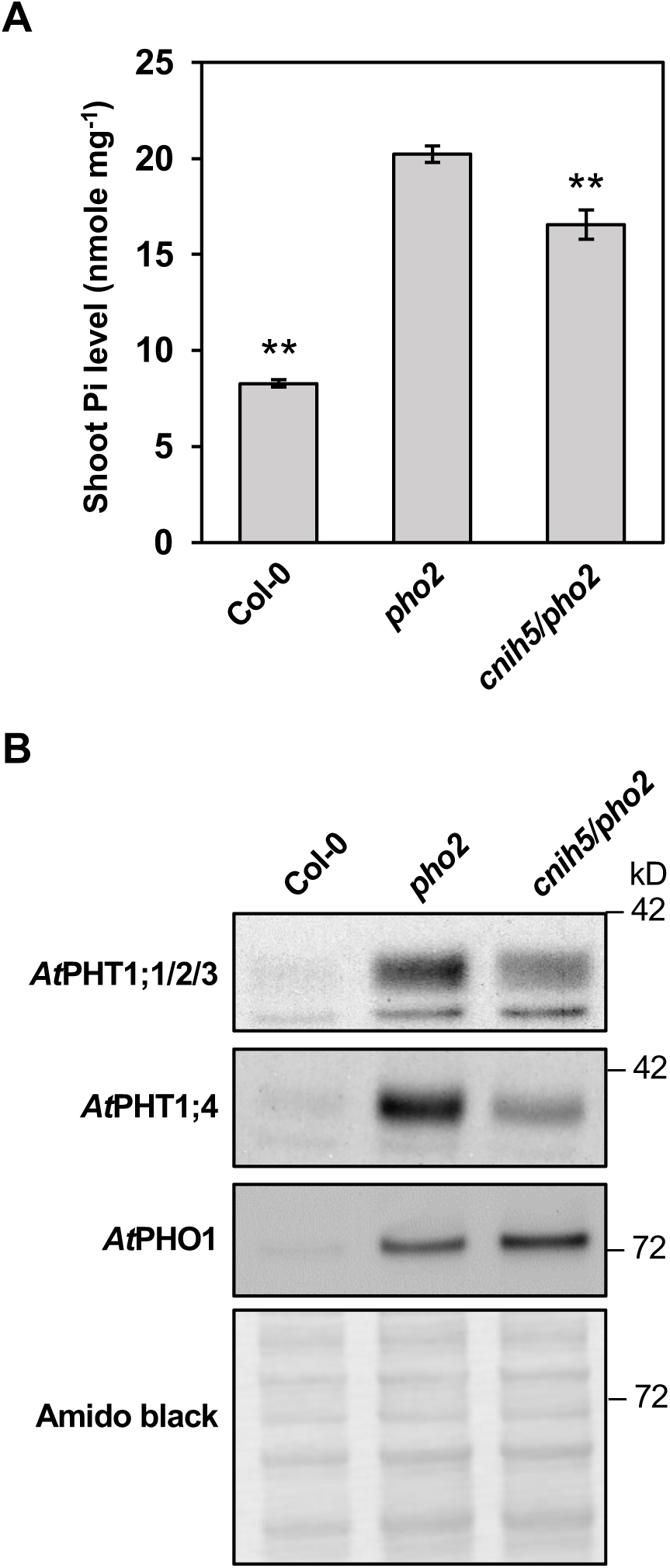
Loss of *AtCNIH5* suppresses Pi accumulation in *Arabidopsis pho2*. (A, B) The shoot Pi level (A) and the expression of *At*PHT1;1/2/3, *At*PHT1;4, and *At*PHO1 in the root (B) of 11-day-old *Arabidopsis* WT, *pho2,* and *cnih5/pho2* seedlings grown under +P (250 µM KH_2_PO_4_) conditions. Error bars represent SE (n = 5–7, pools of seedlings from independent experiments One-way ANOVA with post-hoc Dunnett’s test versus *pho2,* \*\**p* < 0.01. One representative immunoblot from two independent experiments is shown.

## Discussion

### *At*CNIH5 is a Pi starvation-induced and cell type-specific ER cargo receptor for *At*PHT1s

Our study provides multiple lines of evidence showing that *At*CNIH5 is a Pi starvation-induced ER cargo receptor that facilitates the PM targeting of *At*PHT1s. Gene expression profiles using GUS and GFP reporters and subcellular distribution analyses of fluorescent protein fusions suggest *At*CNIH5 as the sole *At*CNIH member induced by low Pi in the root outer layers (**Fig. 1A**, **1B**, and Fig. S2A, S2B, and S3). We also showed that *At*CNIH5 localized to the ER and was closely associated with the ERES markers *At*SAR1A, *At*SEC16A, and *At*SEC24A (**Fig. 2A–C**). As *At*CNIH5 interacts with nearly all the *At*PHT1 family members (**Fig. 3A–C**) (Liu *et al*., 2024), *At*CNIH5 likely participates in the COPII-mediated ER-to-Golgi transport of all *At*PHT1s. We surmise that the recognition of *At*PHT1s by *At*SEC24s alone is not an efficient signal to drive the ER exit of *At*PHT1s, and therefore requires the coincident or successive interaction of *At*PHT1s with *At*CNIH5 and *At*PHF1 (see the discussion in the next section). Although *AtCNIH5* is expressed at a slightly lower level than *AtCNIH1*, *AtCNIH3,* and *AtCNIH4* under Pi sufficiency (Liu *et al*., 2016), the impairment of *AtCNIH5* diminished the amount of *At*PHT1;1–1;4 by 40–50% in the root microsomal fractions (**Fig**. **4C**). The results of confocal imaging also corroborated a lower efficiency of PM targeting and higher intracellular retention of *At*PHT1;1-S11 in the epidermis within the root transition/elongation zone and the root hair of *cnih5* (**Fig. 5A–C**, **Table 1**). Our recent side-by-side membrane proteomic analysis identified *At*PHT1s as the prominent membrane cargoes of *At*CNIH5 under Pi starvation (Liu *et al*., 2024), reinforcing the conclusion that *At*CNIH5 mediates the ER export of *At*PHT1s and, thus, their PM targeting. Since the promoter activities of *AtCNIH1*, *AtCNIH3,* and *AtCNIH4* are barely detected in the epidermal, and cortical, and endodermal cells of the primary root (Fig. S2A), where *AtPHT1;1–1;4* genes are mainly expressed, we infer that *At*CNIH5 is the predominant *At*CNIH member regulating the PM trafficking of *At*PHT1;1–1;4 in these cell layers. However, we cannot exclude the possibility that, in addition to *At*CNIH5, other *At*CNIH members or pairs of *At*CNIHs also mediate the ER export of *At*PHT1 transporters in vascular tissues, even though we did not observe any exacerbated phenotypes of *cnih1/5*, *cnih3/5,* and *cnih4/5* double mutants. Our yeast SUS showed that *At*PHT1;1 also interacts with *At*CNIH1, *At*CNIH3, and *At*CNIH4 (Fig. S10A), implying that different *At*CNIH members could recognize *At*PHT1;1 as their membrane cargo, at least in the yeast heterologous expression system. Of note, we observed a strikingly reduced shoot-to-root distribution of newly taken up Pi in *cnih5* under Pi sufficiency (**Fig. 4D**), indicating that *At*CNIH5 also participates in regulating Pi translocation within plants. This finding aligns with the previously reported but often overlooked role of *At*PHT1;1 in the root-to-shoot translocation of Pi when Pi is abundant (Ayadi *et al*., 2015).

While the *AtCNIH4* transcript is the most abundant in its family according to RNA-seq data (Liu *et al*., 2016) (Fig. S1), the phenotypes of *cnih4* and *cnih4/5* indicate that *At*CNIH4 plays a minor role in modulating Pi homeostasis during the seedling stage. *At*CNIH isoforms have the potential to form tissue– or cell-type-dependent oligomers (Wudick *et al*., 2018), which may affect the ER export of *At*PHT1s to varying degrees. Intriguingly, belonging to the *Brassicaceae* family, which is devoid of mycorrhizal symbiosis, the *Arabidopsis* species was suggested to utilize rapid root growth and root hair formation as a strategy for Pi uptake when grown under Pi depletion (Lambers *et al*., 2008). In agreement with this notion, we observed a less apparent reduction of Pi level in the 11-day-old *cnih5* seedlings (reduction by 18%; **Fig. 4B**) than in the 5-day-old *cnih5* seedlings (reduction by 30%; Fig. S5D), in which macro-scale architectures of the root system were still undeveloped. Based on the PANTHER classification (http://www.pantherdb.org/), *Arabidopsis thaliana* CNIH1 (*At*CNIH1) and *Os*CNIH2 are grouped into the same subfamily as the *yeast* Erv14 and its paralog Erv15; *At*CNIH2, *At*CNIH4, and *Os*CNIH1 belong to the same subfamily; *At*CNIH3 and *At*CNIH5 are *Brassicaceae*-specific CNIH paralogs (Fig. S11). In other words, our data support the notion that the *Brassicaceae*-specific *At*CNIH5 may have a unique physiological function and plays a more crucial role in enhancing Pi transport in the elongating root epidermis and root hairs at the early seedling stage. A previous study on functional complementation in yeast showed that *Sc*Erv14, *At*CNIH1, *At*CNIH3, and *At*CNIH4, but not *At*CNIH2 and *At*CNIH5, enhanced the tolerance of the *Δerv14* mutant to NaCl to various degrees (Wudick *et al*., 2018), suggesting cargo specificity among *At*CNIHs. In our recent study, we identified additional potential membrane cargoes of *At*CNIH5 beyond *At*PHT1s (Liu *et al*., 2024). Interestingly, the first transmembrane domain of *At*CNIH5 is sufficient to interact with *At*PHT1;1, whereas the C-terminal acidic residue of *At*CNIH5 is required for interacting with *At*OCT1 but not with *At*PHT1;1 or *At*DTX21 (Liu *et al*., 2024). Therefore, it is likely that *At*CNIH5 interacts with *At*PHT1s using a mechanism distinct from those used by the acidic domain of fungal and rice CNIHs for their cognate cargoes.

### The interplay of *At*PHF1 and *At*CNIHs in regulating the ER exit of *At*PHT1s

*AtPHF1* is mainly expressed in the root tip and stele under Pi sufficiency and can be further induced in the external root layers at low Pi availability (Bayle *et al*., 2011). This pattern coincides with the Pi starvation-induced expression of *AtCNIH5* in the epidermis and cortex, whereas the gene activity of *AtCNIH5* is absent in the root apical meristem (**Fig. 1B**, Fig. S3). Although the compensation mechanism underlying the upregulation of *At*PHF1 in *cnih5* remains obscure (Fig. S6), the additional loss of *AtCNIH5* in the *phf1* did not synergistically reduce cellular Pi levels but severed plant growth (**Fig. 6B**). This led to the idea that *At*PHF1 and *At*CNIH5 may not play a functionally redundant role but *At*PHF1 is epistatic to *At*CNIH5 in regulating the ER exit of *At*PHT1s. The SEC12-related plant-specific PHF1 was previously proposed to functionally resemble the yeast ER-resident protein Pho86 (González *et al*., 2005). As the *in vitro* vesicle budding assay indicated, the yeast Pho86 may assume a conformation of Pi transporter Pho84 competent for specific incorporation into COPII vesicles or to recruit COPII proteins to the ER (Lau *et al*., 2000). By comparison, the yeast Erv14 cycles between the ER and the Golgi, and the ER membranes lacking functional Erv14 exhibit less efficient incorporation of cargoes into COPII vesicles (Powers & Barlowe, 2002). In accordance, the live-cell imaging results revealed a reduction in the kinetics of ER exit of Erv14’s cargoes in the *erv14Δ* mutant (Herzig *et al*., 2012). Their findings align with our current view on the role of *At*CNIH5 in promoting the ER export of *At*PHT1s. While split-GFP-tagged *At*PHT1;1 was mainly distributed at the PM of root cells in the WT under Pi depletion, it was in part retained intracellularly in the epidermis within the root transition/elongation zone and the root hair of *cnih5* (**Fig. 5B**–**C**, **Table 1**, and Fig. S12). These observations suggested that PM targeting of *At*PHT1;1 is less efficient in the absence of *At*CNIH5. By contrast, *At*PHT1;1-S11 in the *phf1* was barely detectable at the PM of the epidermal cells throughout the different developmental zones, including the lateral root cap (Fig. S12). Unlike the unicellular yeast, we envisage that higher plants have adopted a mechanism to jointly engage PHF1 and CNIHs in specific cell types, thereby selecting and packaging PHT1s for efficient COPII-mediated transport. Indeed, our recent study unveiled that *At*PHF1 interacts with *At*SAR1 GTPase, a key protein that initiates the assembly of COPII, and thus likely participates in the early step of COPII recruitment for the ER export of PHT1s (Lung *et al*., 2025). By contrast, *At*CNIH5 impacts the efficiency of PM trafficking of *At*PHT1s, particularly in the elongating root epidermis and root hairs (**Fig. 5A-C**). As the proliferation of root hair is a crucial trait for improving Pi acquisition under Pi limitation (Gahoonia & Nielsen, 1998, Tanaka *et al*., 2014), it would be interesting to investigate the physiological relevance of *At*CNIH5 in modulating the PM trafficking of *At*PHT1s in root hair cells. It is also worth mentioning that in the yeast SUS, *At*PHF1 interacts with *At*CNIH1, *At*CNIH3, and *At*CNIH4 much less strongly than with *At*CNIH5 (Fig. S10B). This implied that even in the vascular tissues, where *AtPHF1* and *AtCNIH* genes are highly expressed, *At*PHF1 may preferentially interact with *At*CNIH5 over the other *At*CNIH isoforms. According to single-cell mRNA sequencing (scRNA-seq) analyses under high Pi conditions, *AtCNIH1* is mainly expressed in the proliferation and transition domains, with a higher expression level in the phloem of the proliferation domain. *AtCNIH3* is also primarily expressed in the proliferation and transition domains, yet at a lower level, and its expression gradually decreases along the developmental axis as cells differentiate. *AtCNIH4* is expressed in all cell types of the proliferation and transition domains, and strongly expressed in the epidermis of the elongation and maturation domains. *AtCNIH4* and *AtCNIH5* are the most abundant members in epidermal cells among the family. In addition, *AtCNIH5* shows a trend of gradually increased expression in the root epidermis and cortex along the developmental axis and high expression in the stele of the maturation zone (Fig. S13) (Shahan *et al*., 2022, Trevor *et al*., 2022). This highlights a significant role for *At*CNIH5 in facilitating Pi uptake and Pi translocation beyond the root apical meristem, primarily in the outer layers of the non-proliferating root cells upon Pi starvation. Nonetheless, we cannot exclude the possibility that the lack of *At*PHF1 and *At*CNIH5 further suppressed the plant growth due to exaggerated cytoplasmic Pi inadequacy. Alternatively, but not mutually exclusive to this scenario, *At*CNIH5 may have some downstream effectors other than *At*PHT1s that exacerbate the growth retardation of *phf1/cnih5* (Liu *et al*., 2024). On the basis of our present study, we propose a working model as to how *At*PHF1 and *At*CNIH5 interplay to assist the ER exit of *At*PHT1s (**Fig. 8**). *At*PHF1 interacts with *At*PHT1s at the ER to ensure their competence for the ER exit, and likely with *At*SAR1 GTPase as well, accompanied by the COPII assembly at the ERES of nearly every cell type in the root. On the other hand, *At*CNIH5 acts at a later step, perhaps by selectively loading *At*PHT1s at ERES into the COPII-mediated trafficking pathway in the non-dividing root cells, including the epidermis within the transition/elongation zone and the elongating root hair of the differentiation zone. More studies are needed to define the spatiotemporal interaction of *At*PHT1s, *At*PHF1, and *At*CNIH5.

**Fig. 8.**
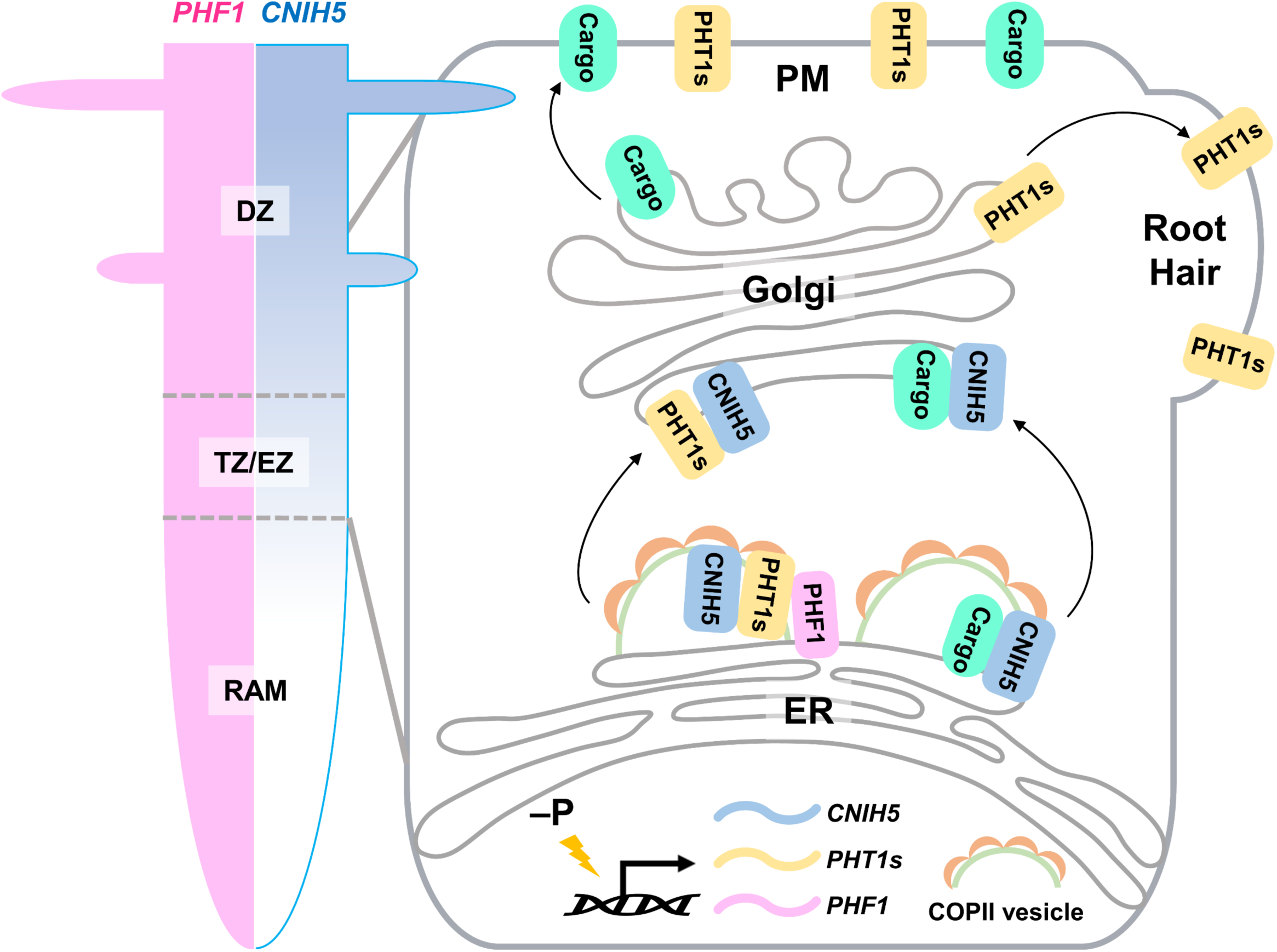
A working model of *At*CNIH5-mediated ER exit of *At*PHT1s. In *Arabidopsi*s, *PHF1* and *CNIH5* are expressed at distinct and overlapping root developmental zones (left panel). In the non-dividing epidermal cell (right panel), including the transition zone/elongation zone and the root hair cell of the differentiation zone, the transcripts of *PHT1* genes, *PHF1*, and *CNIH5* are increased under Pi starvation (−P). CNIH5 interplays with PHF1 to facilitate the ER exit and the plasma membrane (PM) targeting of PHT1s in a COPII-dependent manner. While PHF1 may interact with PHT1s that are competent for the ER exit at an upstream step of COPII recruitment, CNIH5 may selectively mediate the efficient incorporation of PHT1s into the COPII vesicle delivered to the Golgi. CNIH5 may also act as a cargo receptor for other membrane proteins. RAM, root apical meristem; TZ, transition zone; EZ, elongation zone; DZ, differentiation zone.

## Materials and Methods

### Plant material and growth conditions

All *A. thaliana* plants used in this study were in the Columbia-0 (Col-0) ecotype. Seeds of wild-type (WT), *cnih4* (SALK_145991), *cnih5* (SAIL_256_B03), *phf1* (SALK_037068), and *pho2-1* (Delhaize & Randall, 1995) were obtained from the Arabidopsis Biological Resource Center (ABRC). The *cnih1/5* (SAIL_117_E12 for *cnih1* and SAIL_256_B03 for *cnih5*) and *cnih3/5* (GABI_300F09 for *cnih3* and SAIL_256_B03 for *cnih5*) mutants were genotyped as previously described (Wudick *et al*., 2018). *A. thaliana* seeds were surface-sterilized, germinated, and grown on plates with modified one-half strength (1/2) Hoagland nutrient solution containing 250 μM KH_2_PO_4_, 1% sucrose, and 0.8 or 1.2% Bacto agar (BD Difco™ 214010) in the growth chamber at 22℃ with a 16-h light/8-h dark cycle. The Pi-sufficient (+P, 250 μM KH_2_PO_4_) and Pi-deficient (−P, 0 µM KH_2_PO_4_) media were modified 1/2 Hoagland nutrient. The N-deficient (−N, 0 µM and 0.1 mM KNO_3_) media were modified 1/2 Hoagland nutrient solutions. For hydroponic growth of *A. thaliana* adult plants, 1–2-week-old seedlings were transferred to +P (250 μM KH_2_PO_4_) modified 1/2 Hoagland solutions without a supplement of sucrose and grown for additional days in +P (250 μM KH_2_PO_4_) or −P (10 µM KH_2_PO_4_) media as indicated in figure legends. For *Nicotiana benthamiana* (*N. benthamiana*) plants, seeds were surface-sterilized and germinated on 1/2 Murashige and Skoog (MS) medium. Three-to four-week-old plants were used for infiltration of *Agrobacterium tumefaciens* (*A. tumefaciens*).

### Genomic DNA extraction and genotyping of mutants

Genomic DNA was extracted from the leaves of 2-week-old plants with the extraction buffer containing 200 mM Tris-HCl (pH 7.5), 250 mM NaCl, 25 mM EDTA (pH 8.0), and 0.5% SDS. Polymerase chain reaction (PCR) was then performed to validate the presence of the T-DNA alleles of *phf1*, *cnih4*, *cnih5*, and the *pho2* point mutation in samples as previously described. Primer sequences used are listed in Table S1.

### RNA isolation, reverse transcription PCR (RT-PCR), and quantitative RT-PCR

Total root RNA was isolated using GENEzol™ TriRNA Pure Kit with DNase (Geneaid), and a 500-ng RNA was used for the first-strand cDNA synthesis using PrimeScript™ 1st strand cDNA Synthesis Kit (TaKaRa) with oligo(dT) primer. Quantitative RT-PCR (RT-qPCR) was performed using KAPA SYBR® FAST qPCR Master Mix (2×) Kit (Kapa Biosystems) on StepOnePlus™ Real-Time PCR System (Applied Biosystems) according to the manufacturer’s instructions. Gene expression values were normalized to the internal reference *ACT8* (AT1G49240). Transcript expression levels of *AtCNIH* genes in WT, *cnih* single, and *cnih* double mutants were examined by RT-PCR or RT-qPCR. Primer sequences used are listed in Table S1.

### Construct design and yeast split-ubiquitin assay

For constructing the yeast expression clones, the coding sequence (CDS) of *AtCNIH1, AtCNIH3, AtCNIH4, AtCNIH5* and *AtPHT1;4* was amplified by PCR, cloned into the pCR8/GW/TOPO vectors, and sequenced. The pCR8/GW/TOPO constructs containing the CDS of *AtPHT1;1* and *AtPHF1* were cloned as previously reported (Huang *et al*., 2013). The CDS of *AtCNIH5* was recombined into the synthesized MET17-PLV-Cub_GW through an LR reaction, resulting in the N-terminally fused PLVCub-*At*CNIH5. The CDS of *AtPHT1;1* and *At*PHF1 was recombined into MetYC_GW (ABRC plasmid CD3-1740; Obrdlik *et al*., 2004), resulting in the C-terminally fused CubPLV fragments. The CDS of *AtCNIH1, AtCNIH3, AtCNIH4, AtCNIH5*, *AtPHF1*, *AtPHT1;1*, and *AtPHT1;4* was recombined into NX32_GW (ABRC plasmid CD3-1737) through LR reactions, resulting in the N-terminally fused Nub fragments (Obrdlik et al., 2004). The Nub and Cub expression clones were sequentially transformed into the yeast THY.AP4 strain (ABRC CD3-808; Obrdlik et al., 2004). Fresh yeast colonies picked from Yeast Extract Peptone Dextrose Adenine (YPDA) medium or Synthetic Complete (SC) medium without leucine (SC–L) were inoculated in the corresponding liquid medium and incubated overnight at 30℃ with shaking at 200 rpm. Overnight yeast culture was then diluted to OD_600_ = 0.2 and grown for another 5–12 h until OD_600_ reached 0.8. Yeast pellet was collected by centrifugation at 2,000×g at 4℃ for 5 min, washed with ddH_2_O, and resuspended in Tris-EDTA/lithium acetate solution (TE/LiAc, 10 mM Tris-HCl [pH 8.0], 1 mM EDTA, and 0.1 M lithium acetate). The PLVCub*-At*CNIH*5*, *At*PHT1;1– or *At*PHF1-CubPLV-containing transformants were selected and maintained on SC–L for sequential transformation with NubI (positive control), NubG (negative control), NubG-*At*PHT1;1, NubG-*At*PHT1;4, NubG-*At*PHF1, NubG-*At*CNIH1, NubG-*At*CNIH3, NubG-*At*CNIH4, or NubG-*At*CNIH5. Yeast transformants with Nub– and PLVCub-fusions were selected and maintained on SC medium without leucine and tryptophan (SC–LW). For the protein-protein interaction testing, yeast cells co-expressing Nub– and PLVCub-fusions were plated on synthetic media lacking leucine, tryptophan, adenine, and histidine (SC–LWAH) with 0.5 μM or 500 μM methionine or on SC–LW containing 2 g/L 5-bromo-4-chloro-3-indolyl-β-d-galactopyranoside (X-Gal). Primer sequences used are listed in Table S2.

### Construct design for subcellular localization analysis in *N. benthamiana* leaves

For constructing the *UBQ10*:*sXVE*: S10*-At*CNIH5 and *UBQ10*:*sXVE*: mCherry-*At*CNIH5 clones, the CDS of *AtCNIH5* was amplified by PCR and subcloned into *UBQ10*:*sXVE*: S10-(MCS) (Addgene plasmid #108177; Liu et al., 2018, Liu, 2021). For constructing the *UBQ10*: *At*SEC16A-S11 clones, the CDS of *AtSEC16A* was amplified by PCR and subcloned into *UBQ10:* (MCS)-S11. The plasmid encoding *35S*: RFP-*At*CNIH5 was obtained as previously reported (Wudick *et al*., 2018). The plasmid encoding *35S*: ER-mCherry (CD3-959; Nelson *et al*., 2007) was obtained from ABRC. Primer sequences used are listed in Table S2.

### Construct design for the tripartite split-GFP complementation assay in *N. benthamiana* leaves

For constructing the *UBQ10*:*sXVE*: S10-*At*PHF1 and *UBQ10*:*sXVE*: S10-*At*PHO1 clones, the CDS of *AtPHF1* and *AtPHO1* were amplified by PCR and subcloned into *UBQ10*:*sXVE*: S10-(MCS) (Addgene plasmid #108177; Liu et al., 2018, Liu, 2021). For constructing the *UBQ10*:*sXVE*: *At*PHT1;1-S11 and *UBQ10*:*sXVE*: *At*PHO2^C748A^-S11 clones, the CDS of *AtPHT1;1* and *AtPHO2* ^C748A^ were amplified by PCR and subcloned into *UBQ10*:*sXVE*: (MCS)-S11 (Addgene plasmid #108179; (Liu et al., 2018, Liu, 2021). For the tripartite split-GFP complementation, the constructs encoding S10-tagged and S11-tagged protein fusions were co-expressed in the presence of *UBQ10*:*sXVE*: GFP1–9 (Addgene plasmid #108187; Liu et al., 2018, Liu, 2021). Co-expression with S10-3xHA or 3xHA-S11 was a negative control for protein–protein interaction. Primer sequences used are listed in Table S2.

### Construct design and protein degradation assay in *N. benthamiana* leaves

For constructing *UBQ10*:*sXVE*: S10-*At*PHO2 clone, the CDS of *AtPHO2* was amplified and subcloned into pGPTVII. *A. tumefaciens* carrying *UBQ10*:*sXVE*: S10-*At*PHO2, *UBQ10*:*sXVE*: S10-*At*CNIH5 and *UBQ10*:*sXVE*: S10-*At*PHO1 were infiltrated into *N. benthamiana* leaves, each with a final OD_600_ in the range of 0.01–0.05. *A. tumefaciens* carrying *UBQ10*:*sXVE*: GFP (Schlücking *et al*., 2013) was co-infiltrated as an internal control at a final OD_600_ of 0.01. Primer sequences used are listed in Table S2.

### *A. tumefaciens*-mediated infiltration of *N. benthamiana* leaves

*A. tumefaciens* strain EHA105 harboring the expression constructs or p19 was incubated in Luria–Bertani (LB) medium with the appropriate antibiotics [gentamicin (50 μg ml^−1^), rifampicin (5 μg ml^−1^), or kanamycin (50 μg ml^−1^)]. Cells were resuspended in the infiltration medium containing 10 mM MgCl_2_, 10 mM MES (pH 7.5), and 150 μM acetosyringone, and co-infiltrated into the true leaves of 3–4-week-old *N. benthamiana* plants. 36.7 μM β-estradiol was applied for the inducible gene expression as previously described (Liu *et al*., 2018, Liu, 2021). The infiltrated leaves were collected 2 days post-infiltration (dpi) for confocal imaging or protein extraction.

### Construct design and the generation of *A. thaliana* transgenic plants

The CDS of a gene was amplified by PCR, cloned into the pJET1.2/blunt cloning vector, and sequenced before subcloning into the desired expression vectors. For constructing the *AtCNIH1/3/4/5pro:* GUS and *AtCNIH5pro:* GFP reporter clones (Fig. S14), the genomic sequence containing the promoter region of *AtCNIH1* (−497 to +134; transcription start site designated as +1), *AtCNIH3* (−1,793 to +225), *AtCNIH4* (−1,717 to +303), and *AtCNIH5* (−1,701 to +304) was each amplified by PCR and subcloned into pMDC163 or pMDC111.

For constructing the *AtCNIH5* genomic clone (*AtCNIH5pro*: *AtCNIH5*) to complement *cnih5*, the genomic sequence *AtCNIH5*, including the putative promoter region with 5’ UTR (−1,963 to +304) and the genomic sequence containing 3’ UTR (+305 to +1,324), was amplified and cloned into pMDC32 using In-Fusion HD cloning (TaKaRa) for sequencing. Based on the *AtCNIH5* genomic construct clone, the genomic GFP-*AtCNIH5* construct (*AtCNIH5pro*: GFP-*AtCNIH5*) was generated by inserting GFP at the N-terminus of *AtCNIH5* using In-Fusion HD cloning to complement *cnih5*.

For the subcellular localization analysis of *At*PHT1;1-S11, a homozygous line expressing *35S*: GFP1–10 was established and then transformed with the *AtPHT1;1pro*: *At*PHT1;1-S11 construct. For constructing the *35S*: GFP1–10 clone, the CDS of GFP1–10 was amplified and subcloned into pMDC201. For constructing the *AtPHT1;1pro*: *At*PHT1;1-S11 clone, the putative *AtPHT1;1* promoter sequence (−1,964 to +494) and the CDS of *AtPHT1;1* were subcloned into the pGPTVII binary vector. Primer sequences used are listed in Table S2.

### A. tumefaciens-mediated Arabidopsis transformation

Binary expression vectors were transformed into *A. tumefaciens* strain GV3101 (pMP90) and then transformed into *A. thaliana* plants by the floral dip method (Clough & Bent, 1998). T2 transgenic lines with a segregation ratio of 3 resistant: 1 sensitive were selected for T3 propagation. Only T3 homozygous lines were used for further analyses.

### Confocal microscopy

Confocal microscopy images were acquired using Zeiss LSM 800 (Zeiss) with objectives Plan-Apochromat 10×/0.45 M27, 20×/0.8 M27, and 40×/1.3 Oil DIC M27 in multi-track mode with line switching and an averaging of two readings. Additionally, the AiryScan mode was used to enhance the acquisition of punctate structures of GFP-*AtCNIH5* in *AtCNIH5pro*: GFP-*AtCNIH5/cnih5* plants. Excitation/emission wavelengths were 488 nm/410–546 nm for GFP, 561 nm/560–617 nm for RFP and mCherry, and 488 nm/571–700 nm for FM4-64. For subcellular localization analysis of *At*PHT1;1-S11, the seedlings were kept in the dark for 4 h before imaging. All the fluorescence images are acquired from a single confocal optical slice.

### Line profile co-localization analysis

For high-resolution co-localization of *35S:* RFP-*At*CNIH5 or *UBQ10*:*sXVE*: mCherry-*At*CNIH5 with the *cis*-Golgi marker MNS1 and the three ERES markers *At*SAR1A, *At*SEC16A, and *At*SEC24A in *N. benthamiana* leaves, images were acquired using the line profile utility in Zeiss ZEN Blue software as previously reported (McGinness *et al*., 2025). Only well-oriented side-profile views that allowed accurate visualization of cisternal separation were used for quantitative co-localization analysis. For each suitable Golgi body, a line was drawn perpendicular to the organelle’s long axis to generate fluorescence intensity profiles for the two channels. The distance between the maximum intensity peaks of each channel was measured across the line profile, providing a quantitative estimate of spatial separation between fluorescent signals. Data extraction and processing were automated using a Python (v3.9) script (McGinness et al., 2025). The script generated both Gaussian-fitted and raw distance datasets to verify accuracy and consistency, from which average inter-peak distances and associated standard deviations were computed. Kruskal–Wallis analysis was used to compare the median peak distances among multiple groups. Post-hoc pairwise comparisons were performed using Dunn’s multiple comparison test with Bonferroni correction to identify specific group differences.

### Histochemical staining of β-glucuronidase (GUS)

GUS staining was performed as previously described (Li, 2011). Seedlings were submerged in freshly prepared X-Gluc staining buffer containing 500 mM NaH_2_PO_4_, 500 mM Na_2_HPO_4_ · 7H_2_O, 1mM K_3_Fe(CN)_6_, 1 mM K_4_Fe(CN)_6_, 10 mM EDTA, 0.1% Triton X-100, and 2.25 mM X-glucuronide (Cyrusbioscience), vacuumed for 20 min, and then incubated at 37℃ for 2–2.5 h. The pigments were washed out serially using 10–99% ethanol for 30 min. Finally, the GUS staining was examined using a stereo microscope (Prompt Ocean ST3E-0745 or Leica DM2500).

### Pi concentration and Pi uptake assays

Pi concentrations were analysed as described (Ames, 1966) with minor modifications. For the Pi assay, fresh tissue was homogenized with 1% glacial acetic acid and incubated at 42℃ for 30 min, followed by centrifugation at 13,000×g for 5 min to obtain a clear supernatant. The supernatant aliquot was mixed with the assay solution (0.35% NH_4_MoO_4_, 0.86 N H_2_SO_4,_ and 1.4% ascorbic acid) and incubated at 42℃ for an additional 30 min before measurement at A_750_. The Pi uptake assay for whole seedlings was conducted in +P (250 µM KH_2_PO_4_) or −P (10 µM KH_2_PO_4_) medium supplemented with ^32^P as previously described (Kanno *et al*., 2016).

### Total protein and microsomal protein extraction

For the extraction of root total proteins, samples were ground in liquid N and dissolved in protein lysis buffer containing 60 mM Tris-HCl (pH 8.5), 2% SDS, 2.5% glycerol, 0.13 mM EDTA, 1 mM phenylmethylsulfonyl fluoride (PMSF), and Protease Inhibitor Cocktail (Sigma-Aldrich). Root microsomal proteins were extracted using the Minute™ Plant Microsomal Membrane Extraction Kit (Invent Biotechnologies) and were resuspended in the same protein lysis buffer that was used for total protein extraction.

### Immunoblot analysis

For *Arabidopsis* seedlings, a total of 25 μg of root total proteins or 15 μg of root microsomal proteins from each sample were mixed with LDS sampling buffer containing 1 mM DTT. The mixture was then denatured by heating and loaded onto 4–12% Q-PAGE™ Bis-Tris Precast Gel (SMOBIO) or NuPAGE 4–12% Bis-Tris Gels (Thermo Fisher Scientific). For *N. benthamiana* leaves, 25 μg of total protein was used for loading. Gel was transferred to polyvinylidene difluoride membranes (Immobilon-P Membrane or Immobilon®-PSQ Membrane). Membrane was blocked with 1 or 2% BSA in 1× PBS solution with 0.2% Tween 20 (PBST, pH 7.2) at RT for 1 h, and washed 4 times with 1× PBST for 5 min followed by hybridization with primary antibodies of *At*PHT1;1/2/3 (1:1,000, PHY0753, PhytoAB), *At*PHT1;4 (Huang *et al*., 2013), GFP (1:5,000, AS214696, Agrisera) and S10 (1:1,000) for overnight at 4℃; *At*PHF1 (Huang *et al*., 2013), *At*PHO1 (Liu *et al*., 2012), and *At*CNIH5 (Liu *et al*., 2024) at RT for 2 h. Anti-S10 polyclonal rabbit antibody was raised against the peptide of S10 (MDLPDDHYLSTQTILSKDLN) (Liu *et al*., 2018) and produced by LTK BioLaboratories, Taiwan. Anti-*At*CNIH5 polyclonal rabbit antibody was raised and used as described (Liu *et al*., 2024). The membrane was washed four times with 1× PBST for 5 min, followed by hybridization with the horseradish peroxidase–conjugated secondary antibody (1:20,000, GTX213110-01 and 1:10,000, GTX213111-01, GeneTex) at RT for 1 h. After four washes in 1× PBST for 5 min and a rinse with distilled water, chemiluminescent substrates (WesternBright ECL and WesternBright Sirius ECL, Advansta) were applied for signal detection.

### Co-immunoprecipitation

The root of 11-day-old *Arabidopsis UBQ10*: GFP and *AtCNIH5pro*: GFP-*AtCNIH5/cnih5* complementation (COM) seedlings were grown under –P (0 µM KH_2_PO_4_, seven days of starvation) conditions and ground in liquid nitrogen. The root membrane fraction was isolated using the low-speed pellet (LSP) method as previously described (Yoshimoto *et al*., 2004) and resuspended in solubilization buffer containing 100 mM Tris-HCl (pH 7.5), 400 mM sucrose, 1 mM EDTA, 0.6% or 1% Triton X-100, 1 mM phenylmethylsulfonyl fluoride and Protease Inhibitor Cocktail (Sigma-Aldrich). An equivalent of 30 μg or 100 μg of root membrane-enriched protein was loaded onto 5 μl of GFP-Trap beads (Chromotek) and incubated at 4℃ for 2 h. The agarose beads were then washed once with the abovementioned solubilization buffer containing 0.1% Triton X-100 and resuspended in 1× Laemmli buffer. The mixture was boiled at 95℃ for 5 min. The centrifuged supernatant was collected for immunoblot analysis.

### Root growth analysis

Seedlings were grown on +P (250 µM KH_2_PO_4_) or −P (10 µM KH_2_PO_4_) media. For the measurement of primary and lateral root lengths on media containing 1.2% Bacto agar, photos of roots were taken by a PowerShot G16 Camera (Canon), and the length was measured using ImageJ segmented line function (Schneider *et al*., 2012). Only lateral roots with lengths longer than 2 mm were included for measurement. For the measurement of root hair length, seedlings were grown on media containing 0.6% Bacto agar. Root hairs were observed under the Leica DM2500 microscope equipped with DIC optics and photographed. The ten longest root hairs within the maturation zone per root, as defined from 5 mm to 10 mm away from the root tip, were measured by ImageJ. Root hair numbers within the above-mentioned maturation zone were counted for root hair density.

### Chemical treatments

Acetosyringone (150 mM stock in DMSO; Sigma-Aldrich) and β-estradiol (36.7 mM stock in ethanol; Sigma-Aldrich) were diluted to final concentrations of 150 µM and 36.7 μM, respectively, in ddH_2_O.

### Accession Numbers

Sequence data from this article can be found in the Arabidopsis Genome Initiative under the following accession numbers: *At*CNIH1 (AT3G12180), *At*CNIH3 (AT1G62880), *At*CNIH4 (AT1G12390), *At*CNIH5 (AT4G12090), *At*PHT1;1 (AT5G43350), *At*PHT1;4 (AT2G38940), *At*PHF1 (AT3G52190), *At*PHO1 (AT3G23430), *At*PHO2 (AT2G33770), *At*SEC16A (AT5G47480), *At*SEC24A (AT3G07100), and *At*SAR1A (AT2G33120).

## Funding

This work is supported by the Ministry of Science and Technology (MOST 108-2311-B-007-003-MY3) and the National Science and Technology Council (NSTC 112-2313-B-007-001-MY3).

## Acknowledgements

We thank Prof. José A. Feijó at the University of Maryland, USA, for sharing the *Arabidopsis cnih1/5* and *cnih3/5* mutants and plasmid encoding *35S*: RFP-*At*CNIH5. We thank Dr. Tzyy-Jen Chiou at Academia Sinica, Taiwan (R.O.C.), for sharing the *Arabidopsis phf1* mutants, anti-*At*PHT1;4, anti-*At*PHF1, and anti-*At*PHO1 antibodies. We acknowledge Ms. Wen-Chun Chou for constructing the plasmid encoding *35S*: GFP1–10 and Mr. Po-Ruey Huang for cloning the NubG-*At*PHT1;4 construct. We acknowledge Mr. Hong-Po Chen and Ms. Hui-Ying Chen for the genotyping of *phf1/cnih* mutants and of *AtPHT1;1pro*: *At*PHT1;1-S11*/35S*: GFP1–10 in the WT and *cnih5* backgrounds and the technical support from the confocal imaging core, National Tsing Hua University (sponsored by MOST 108-2731-M-007-001 and MOST 110-2731-M-007-001). We are grateful to Dr. Laurent Nussaume at CEA Cadarache, France, and Dr. Tzyy-Jen Chiou for valuable comments on the manuscript.

## Competing interests

None declared

## Author contributions

T.-Y. L. conceived the project, designed, performed and analysed the experiments. C.-Y. C., C.-D. T., H.-F. L., J.-Y. W., M.-H. T., A. J. M., S. K., V. K. and C.-A. L. performed and analysed the experiments. T.-Y. L. and C.-Y. C. wrote the article.

## Data availability

All data supporting the findings of this study are available within the article and the Supporting information (Figs S1–S14, Tabs S1–S2).

## Supporting information

**Fig. S1.** Expression of *AtCNIH* genes under Pi deprivation by RNA-seq analysis. Expression of *AtCNIH1/2/3/4/5* in the shoot and root of 10-day-old WT (Col-0) seedlings under Pi-sufficient conditions (+P0) or under one day (–P1) and three days of Pi starvation (–P3) as previously described (Liu *et al*., 2016). RPKM stands for reads *per* kilobase of transcript *per* million mapped reads. Numbers represent average RPKM values of two replicates, with an assigned value of 0.25 for readings below this threshold.

**Fig. S2.** Expression of *AtCNIH1pro:* GUS, *AtCNIH3pro:* GUS and *AtCNIH4pro:* GUS in *Arabidopsis*. (A, B) Expression of *AtCNIH1pro*: GUS, *AtCNIH*3*pro*: GUS, and *AtCNIH4pro*: GUS in the primary (A) and lateral roots (B) of 7-day-old *Arabidopsis* seedlings grown under +P (250 µM KH_2_PO_4_ and 7.5 mM KNO_3_), –P (0 µM KH_2_PO_4_ and 7.5 mM KNO_3_, five days of starvation), or –N (250 µM KH_2_PO_4_ and 0.1 mM KNO_3_, five days of starvation) conditions. Representative images are shown from three independent T3 homozygous lines. Scale bars, 100 µm.

**Fig. S3.** Expression of *AtCNIH5pro:* GFP in *Arabidopsis*. Expression of *AtCNIH*5*pro*: GFP in 5-day-old *Arabidopsis* seedlings grown under +P (250 µM KH_2_PO_4_) or –P (0 µM KH_2_PO_4_, five days of starvation) conditions. The maturation zone, basal meristem, and apical meristem of the primary root are shown as indicated. Merged, single confocal sliced images of the green fluorescence and bright-field channels are shown. Representative images are shown from three independent T3 homozygous lines. Scale bars, 50 µm.

**Fig. S4.** Complementation of *cnih5* by the expression of the genomic *AtCNIH5* and GFP-*AtCNIH5* sequences. (A–B) The shoot fresh weight (FW), Pi content, and Pi level of 11-day-old seedlings of *Arabidopsis* WT, *cnih5*, and *AtCNIH5pro*: GFP-*AtCNIH5/cnih5* complementation (COM) lines (A) or *AtCNIH5pro*: *AtCNIH5/cnih5* COM lines (B) grown under +P (250 µM KH_2_PO_4_) conditions. Error bar represents SE (n = 7 or 9 pools of seedings from independent experiments). One-way ANOVA with post-hoc Dunnett’s test versus *cnih5,* \**p* < 0.05, \*\**p* < 0.01. (C) Protein expression of *At*PHT1;1/2/3, *AtCNIH5*, or GFP-*AtCNIH5* in the low-speed pellet (LSP) of *Arabidopsis* WT, *cnih5,* and COM lines under –P (0 µM KH_2_PO_4_, three days of starvation) conditions. The relative expression level was normalized with the corresponding amido black staining and relative to the WT control. One representative immunoblot from two independent experiments is shown, and quantitative data are indicated as mean ± SE (n = 2) for the relative expression of *At*PHT1;1/2/3. Root microsomal protein was isolated by the LSP method as previously described (Yoshimoto *et al*., 2004).

**Fig. S5.** The expression of Pi starvation-induced genes and the root phenotypes of *Arabidopsis cnih5.* (A) RT-qPCR analysis of the transcript expression of *AtPHT1;1–1;4*, *AtACP5*, and *AtPHF1* in the root of 11-day-old *Arabidopsis* WT and *cnih5* seedlings grown under +P (250 µM KH_2_PO_4_) or –P (0 µM KH_2_PO_4_, three days of starvation) conditions. (B, C) The primary root (PR) length (B) and the lateral root (LR) density (C) of 9-day-old *Arabidopsis* WT and *cnih5* seedlings grown under +P (250 µM KH_2_PO_4_) or –P (10 µM KH_2_PO_4_, five days of starvation) conditions. (D*–*F) The shoot Pi level (D), the root hair (RH) length (E), and the RH density (F) of 5-day-old *Arabidopsis* WT and *cnih5* seedlings grown under +P (250 µM KH_2_PO_4_) or –P (10 µM KH_2_PO_4_, three days of starvation in D, E or 0 µM KH_2_PO_4_, five days of starvation in F). Data in (B, C, E, and F) are visualized using BoxPlotR (Spitzer *et al*., 2014). Data points are plotted as dots. The number of dots is indicated at the bottom of each figure. The centre lines show the medians; the central plus signs (+) show the means; box limits indicate the 25th and 75th percentiles; whiskers extend to the minimum and the maximum values. Error bar represents SE in D (n = 3, pools of seedlings from independent experiments). Data significantly different from the corresponding WT in (D) are indicated by asterisks (\**p* < 0.05; Student’s t-test, two-tailed) and from the other groups in (B), (C), (E), and (F) are indicated by different lowercase letters (Two-Way ANOVA, Duncan’s test).

**Fig. S6.** Decreased *At*PHT1s and increased *At*PHF1 in the total root proteins from *Arabidopsis cnih5*. Protein expression of *At*PHT1;1/2/3, *At*PHT1;4 and *At*PHF1 in the root of 11-day-old seedlings of *Arabidopsis* WT, *cnih4,* and *cnih5* under +P (250 µM KH_2_PO_4_) or –P (0 µM KH_2_PO_4_, three days of starvation) conditions. The relative expression level of *At*PHT1;1/2/3, *At*PHT1;4, and *At*PHF1 was normalized with the corresponding actin and relative to the WT control. Error bars represent SE (n = 3–4, pools of seedlings from independent experiments). One-way ANOVA with post-hoc Dunnett’s test versus WT, \**p* < 0.05.

**Fig. S7.** Analyses of *Arabidopsis cnih1/5, cnih3/5*, and *cnih4/5* seedlings. (A) The transcript expression of *AtCNIH* genes in the root of 11-day-old *Arabidopsis* WT, *cnih5*, *cnih1/5*, *cnih3/5*, and *cnih4/5* seedlings grown under +P (250 µM KH_2_PO_4_) conditions assessed by RT-qPCR or RT-PCR. (B, C) The shoot and root fresh weight (FW) (B) and Pi levels (C) of 11-day-old *Arabidopsis* WT, *cnih5*, *cnih1/5*, *cnih3/5*, and *cnih4/5* seedlings grown under +P (250 µM KH_2_PO_4_) or –P (0 µM KH_2_PO_4_, three days of starvation) conditions. Error bars represent SE (n = 6–10, pools of seedlings from independent experiments). One-way ANOVA with post-hoc Dunnett’s test versus *cnih5*, \**p* < 0.05.

**Fig. S8.** Phenotypes of *Arabidopsis phf1, phf1/cnih4, phf1/cnih5*, and *phf1/cnih4/5* adult plants. (A, B) The shoot and root fresh weight (FW) (A) and Pi levels (B) of 20-day-old *Arabidopsis* WT, *phf1*, *phf1/cnih*4, *phf1/cnih5*, and *phf1/cnih4/5* plants grown under +P (250 µM KH_2_PO_4_) or –P (10 µM KH_2_PO_4_, five days of starvation) hydroponic conditions. Data are visualized using BoxPlotR (Spitzer *et al*., 2014). Centre lines show the medians; the central plus signs (+) show the means; box limits indicate the 25th and 75th percentiles; whiskers extend to the 5th and 95th percentiles, and data points are plotted as dots. Data significantly different from the other groups are indicated by different lowercase letters (n = 15–24, biological replicates from independent experiments; One-Way ANOVA, Duncan’s test).

**Fig. S9.** *At*PHO2 does not mediate the degradation of *At*CNIH5. (A) Co-expression of S10-*At*CNIH5 with the catalytically defective *At*PHO2^C748A^-S11 in the presence of GFP1–9 in *N. benthamiana* leaves. Images are from single confocal slices. Scale bars, 20 µm. The co-expression of S10-*At*CNIH5 with *At*PHT1;1-S11 and S10-3xHA-S11 are, respectively, used as positive and negative protein–protein interaction controls. (B) Immunoblot analysis of S10-*At*CNIH5 and S10-*At*PHO1 in *N. benthamiana* leaves under co-expression of increasing levels of S10-*At*PHO2. The co-expression of GFP is used as a negative control. Anti-S10 antibodies also detect free GFP.

**Fig. S10.** Interaction analysis of *At*PHT1;1 and *At*PHF1 with *At*CNIH1, *At*CNIH3, and *At*CNIH4. (A, B) Co-expression of *At*PHT1;1-PLVCub (A) and *At*PHF1-PLVCub (B) with NubG-*At*CNIH1, –*At*CNIH3, or –*At*CNIH4 in the yeast split-ubiquitin system. The co-expression of *At*PHT1;1-PLVCub or *At*PHF1-PLVCub with NubI and NubG are used as positive and negative controls, respectively. Yeast transformants were grown on synthetic medium without leucine and tryptophan (–LW; the left panel) or on synthetic medium lacking leucine, tryptophan, adenine, and histidine-containing 0.5 μM methionine (–LWAH; middle panel) or on SC–LW containing 2 mg/L X-gal (X-gal; the right panel).

**Fig. S11.** The CORNICHON-RELATED family (PTHR12290) in Protein Analysis THrough Evolutionary Relationships (PANTHER). Green circles represent speciation events, while orange circles represent duplication. Diamonds are expanded subfamily nodes, and triangles are collapsed nodes. Note that we have collapsed some nodes (shown with triangles) here to simplify the diagram by hiding some descendant subtrees. Different colors correspond to different subfamilies. The whole family tree can be explored at www.pantherdb.org/treeViewer/treeViewer.jsp?book= PTHR12290

**Fig. S12.** Impaired plasma membrane targeting of *At*PHT1;1-S11 in *Arabidopsis phf1*. Representative images of localization of *At*PHT1;1-S11 in 5-day-old *AtPHT1;1pro*: *At*PHT1;1-S11/*35S*: GFP1–10 seedlings grown under −P (0 μM KH_2_PO_4_) conditions. Independent lines in the *phf1* and *cnih5* backgrounds, and the corresponding WT controls are used for comparison. Arrows indicate GFP signals surrounding the nuclear membrane. RAM, Root apical meristem; TZ, transition zone; EZ, elongation zone; DZ, differentiation zone. Images are from single confocal slices. Scale bar, 20 µm.

**Fig. S13.** Expression of *AtPHT1;1*, *AtPHT1;2*, *AtPHT1;4*, *AtPHT1;5*, *AtPHT1;7–1;9*, *AtCNIH1–5,* and *AtPHF1* revealed by single-cell mRNA sequencing (scRNA-seq) of *Arabidopsis* root. The dot plot used to visualize single-cell gene expression in *Arabidopsis* roots is retrieved from ARVEX (https://shiny.mdc-berlin.de/ARVEX/), which hosts processed scRNA-seq data of wild-type samples from Shahan *et al*. (2022) and Trevor *et al*. (2023). PctExp: Percentage of cells expressing the gene in a specific time zone and cell type; circle size represents the proportion (%) of cells expressing a given gene. AvgExp: Averaged normalized gene expression value in a specific time zone and cell type; the colour of the scale bar represents the mean expression (natural log +1 pseudocount).

**Fig. S14.** Schematic representations of *AtCNIH* gene promoter-fused GUS and GFP reporter constructs. Schematic representations of *AtCNIH1*, *AtCNIH3*, *AtCNIH4*, and *AtCNIH5* promoter-fused GUS reporter constructs (A) and *AtCNIH5* promoter-fused GFP reporter construct (B). The upstream region of transcription started site (TSS) in *AtCNIH* genes was shown by a white arrow, and gray boxes showed the 5’ untranslated regions.

**Table S1.** Oligonucleotides used for PCR genotyping, RT-PCR, and RT-qPCR analyses.

**Table S2.** Oligonucleotides used for plasmid constructs.

## Notes

### Competing Interest Statement

The authors have declared no competing interest.

### Summary of Updates

New results are added, or data are presented in a new format.

