## Supplemental Figures and Table for "Phosphate starvation-induced CORNICHON HOMOLOG 5 as endoplasmic reticulum cargo receptor for PHT1 transporters in *Arabidopsis*"

| AGI | Gene Name | Shoot | | | Root | | |
|---|---|---|---|---|---|---|---|
| | | +P0 | –P1 | –P3 | +P0 | –P1 | –P3 |
| AT3G12180 | *AtCNIH1* | 9.54 | 8.82 | 8.40 | 17.01 | 16.24 | 17.61 |
| AT1G12340 | *AtCNIH2* | 0.29 | 0.25 | 0.25 | 0.25 | 0.25 | 0.25 |
| AT1G62880 | *AtCNIH3* | 4.81 | 4.05 | 3.82 | 10.77 | 10.69 | 10.93 |
| AT1G12390 | *AtCNIH4* | 19.59 | 11.60 | 11.72 | 30.82 | 27.85 | 26.73 |
| AT4G12090 | *AtCNIH5* | 1.49 | 1.41 | 3.61 | 9.59 | 17.35 | 60.62 |

**Fig. S1** Expression of *AtCNIH* genes under Pi deprivation by RNA-seq analysis. Expression of *AtCNIH1/2/3/4/5* in the shoot and root of 10-day-old WT (Col-0) seedlings under Pi-sufficient conditions (+P0) or under one day (–P1) and three days of Pi starvation (–P3) as previously described (Liu *et al*., 2016). RPKM stands for reads *per* kilobase of transcript *per* million mapped reads. Numbers represent average RPKM values of two replicates, with an assigned value of 0.25 for readings below this threshold.

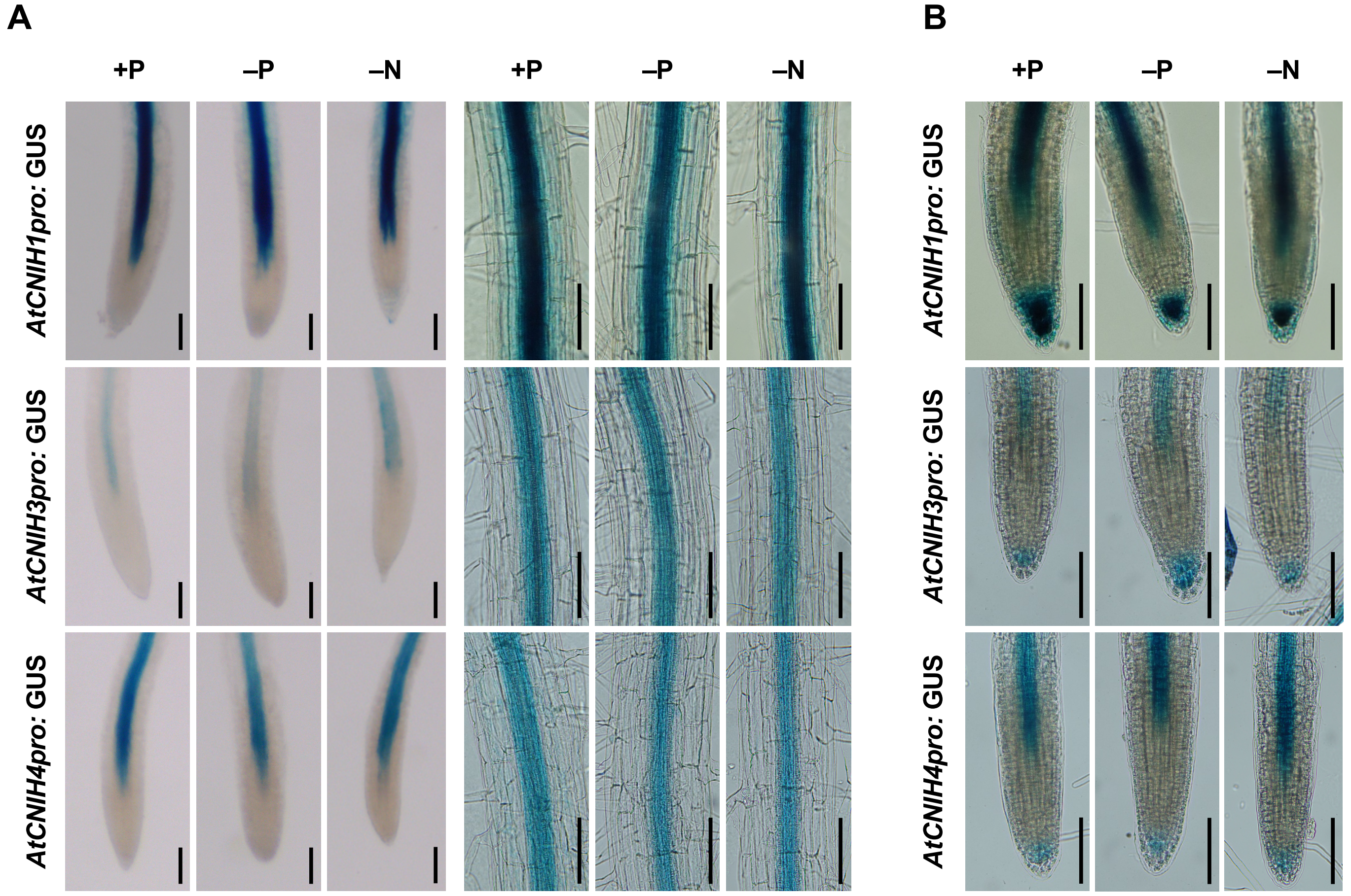


**Fig. S2** Expression of *AtCNIH1pro:* GUS, *AtCNIH3pro:* GUS and *AtCNIH4pro:* GUS in *Arabidopsis*. (A, B) Expression of *AtCNIH1pro*: GUS, *AtCNIH3pro*: GUS, and *AtCNIH4pro*: GUS in the primary (A) and lateral roots (B) of 7-day-old *Arabidopsis* seedlings grown under +P (250 μM $KH_2PO_4$ and 7.5 mM $KNO_3$), –P (0 μM $KH_2PO_4$ and 7.5 mM $KNO_3$, five days of starvation), or –N (250 μM $KH_2PO_4$ and 0.1 mM $KNO_3$, five days of starvation) conditions. Representative images are shown from three independent T3 homozygous lines.Scale bars, 100 μm.

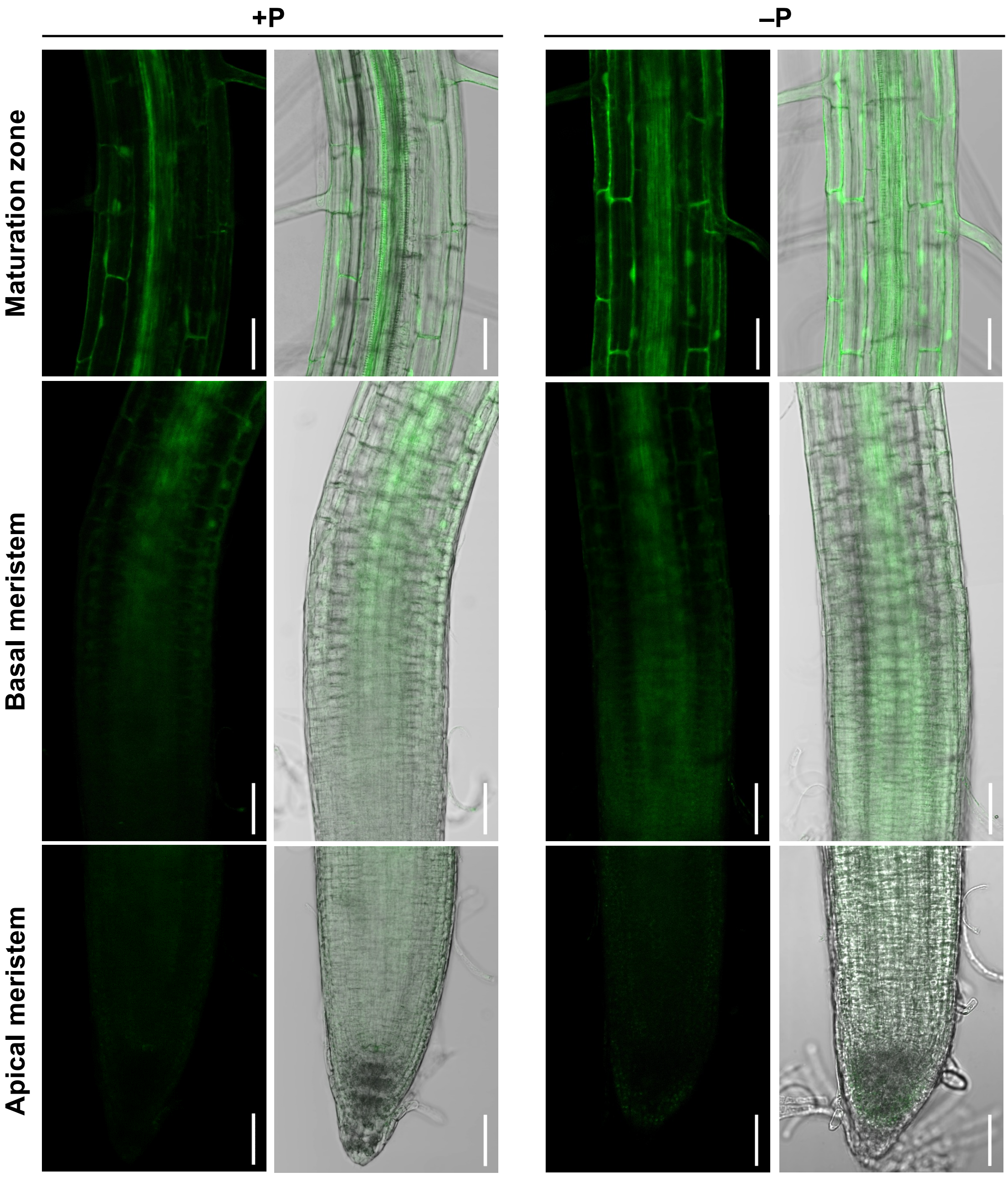


**Fig. S3** Expression of *AtCNIH5pro:* GFP in *Arabidopsis*. Expression of *AtCNIH5pro*: GFP in 5-day-old *Arabidopsis* seedlings grown under +P (250 µM $KH_2PO_4$) or –P (0 µM $KH_2PO_4$, five days of starvation) conditions. The maturation zone, basal meristem, and apical meristem of the primary root are shown as indicated. Merged, single confocal sliced images of the green fluorescence and bright-field channels are shown. Representative images are shown from three independent T3 homozygous lines. Scale bars, 50 µm.

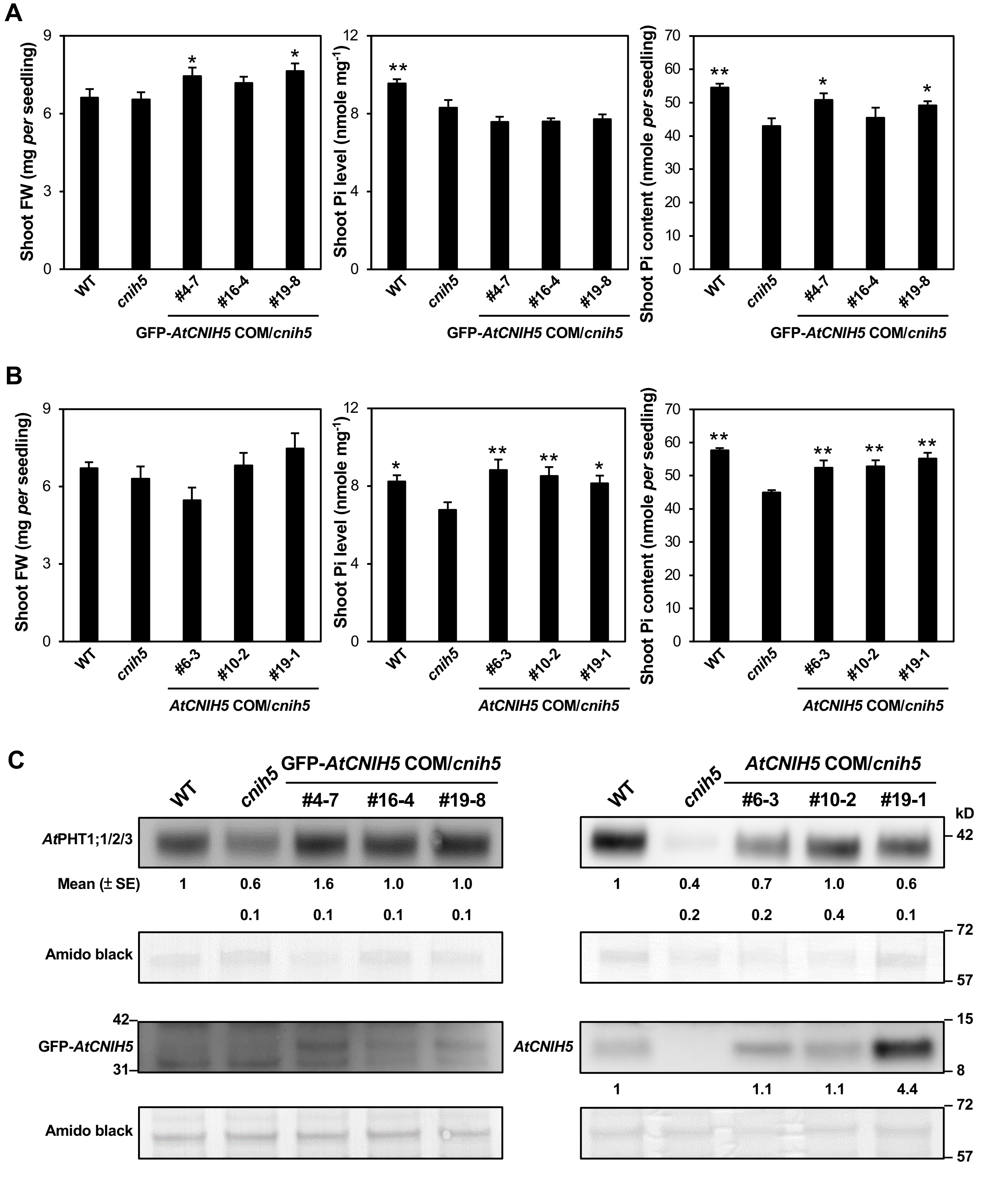


**Fig. S4** Complementation of *cnih5* by the expression of the genomic *AtCNIH5* and GFP-*AtCNIH5* sequences. (A–B) The shoot fresh weight (FW), Pi content, and Pi level of 11-day-old seedlings of *Arabidopsis* WT, *cnih5*, and *AtCNIH5pro*: GFP-*AtCNIH5/cnih5* complementation (COM) lines (A) or *AtCNIH5pro*: *AtCNIH5/cnih5* COM lines (B) grown under +P (250 μM $KH_2PO_4$) conditions. Error bar represents SE (n = 7 or 9 pools of seedings from independent experiments). One-way ANOVA with post-hoc Dunnett's test versus *cnih5*, $^{*}p < 0.05$, $^{**}p < 0.01$. (C) Protein expression of *At*PHT1;1/2/3, *AtCNIH5*, or GFP-*AtCNIH5* in the low-speed pellet (LSP) of *Arabidopsis* WT, *cnih5,* and COM lines under –P (0 μM $KH_2PO_4$, three days of starvation) conditions. The relative expression level was normalized with the corresponding amido black staining and relative to the WT control. One representative immunoblot from two independent experiments is shown, and quantitative data are indicated as mean ± SE (n = 2) for the relative expression of *At*PHT1;1/2/3. Root microsomal protein was isolated by the LSP method as previously described (Yoshimoto *et al*., 2004).

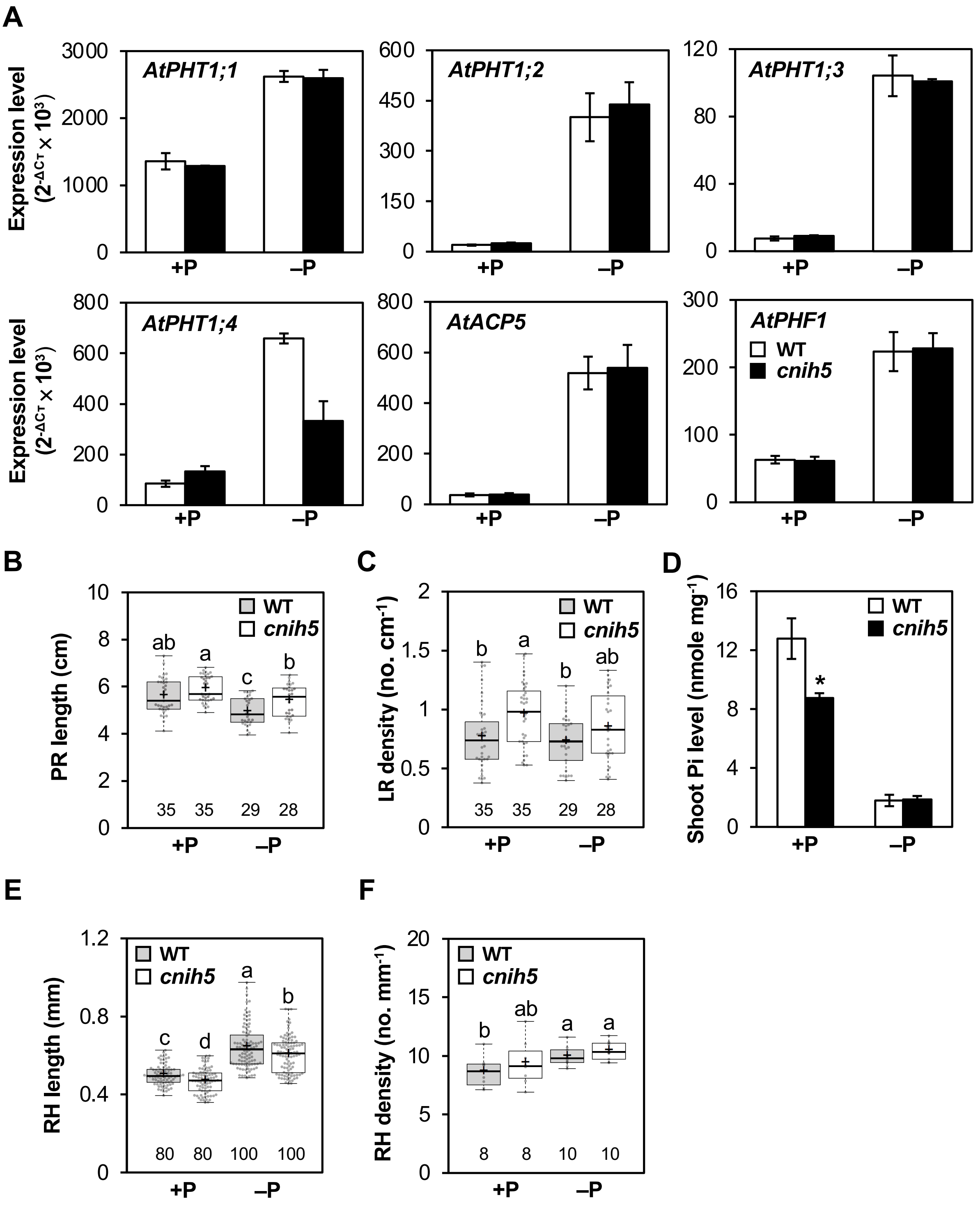


**Fig. S5** The expression of Pi starvation-induced genes and the root phenotypes of *Arabidopsis cnih5.* (A) RT-qPCR analysis of the transcript expression of *AtPHT1;1–1;4*, *AtACP5*, and *AtPHF1* in the root of 11-day-old *Arabidopsis* WT and *cnih5* seedlings grown under +P (250 μM $KH_2PO_4$) or –P (0 μM $KH_2PO_4$, three days of starvation) conditions. (B, C) The primary root (PR) length (B) and the lateral root (LR) density (C) of 9-day-old *Arabidopsis* WT and *cnih5* seedlings grown under +P (250 μM $KH_2PO_4$) or –P (10 μM $KH_2PO_4$, five days of starvation) conditions. (D–F) The shoot Pi level (D), the root hair (RH) length (E), and the RH density (F) of 5-day-old *Arabidopsis* WT and *cnih5* seedlings grown under +P (250 μM $KH_2PO_4$) or –P (10 μM $KH_2PO_4$, three days of starvation in D, E or 0 μM $KH_2PO_4$, five days of starvation in F). Data in (B, C, E, and F) are visualized using BoxPlotR (Spitzer *et al.*, 2014). Data points are plotted as dots. The number of dots is indicated at the bottom of each figure. The centre lines show the medians; the central plus signs (+) show the means; box limits indicate the 25th and 75th percentiles; whiskers extend to the minimum and the maximum values. Error bar represents SE in D (n = 3, pools of seedlings from independent experiments). Data significantly different from the corresponding WT in D are indicated by asterisks (*$p < 0.05$; Student's t-test, two-tailed) and from the other groups in B, C, E, and F are indicated by different lowercase letters (Two-Way ANOVA, Duncan's test).

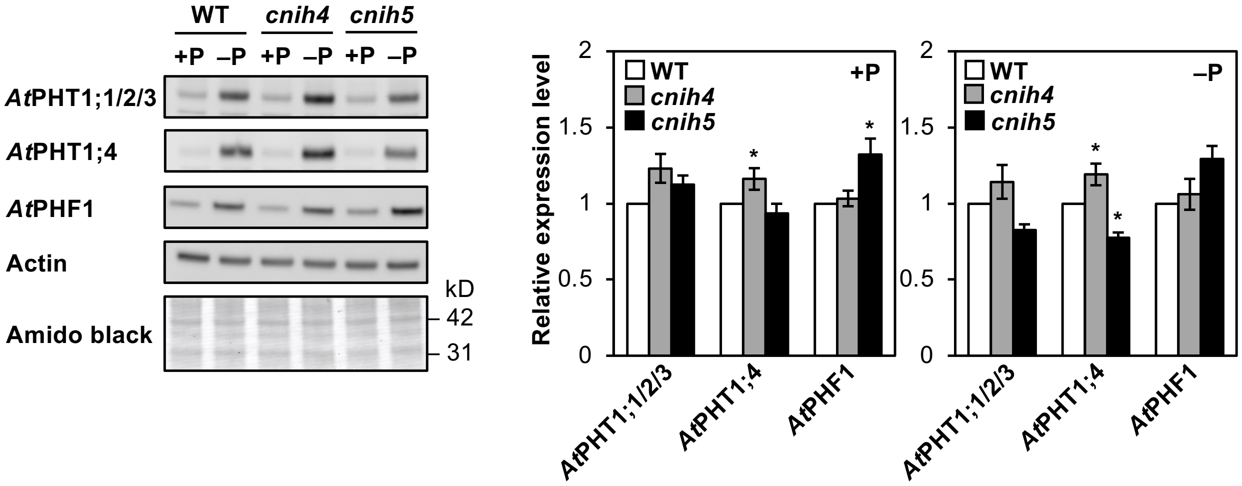


**Fig. S6** Decreased *At*PHT1s and increased *At*PHF1 in the total root proteins from *Arabidopsis cnih5*. Protein expression of *At*PHT1;1/2/3, *At*PHT1;4 and *At*PHF1 in the root of 11-day-old seedlings of *Arabidopsis* WT, *cnih4,* and *cnih5* under +P (250 µM $KH_2PO_4$) or –P (0 µM $KH_2PO_4$, three days of starvation) conditions. The relative expression level of *At*PHT1;1/2/3, *At*PHT1;4, and *At*PHF1 was normalized with the corresponding actin and relative to the WT control. Error bars represent SE (n = 3–4, pools of seedlings from independent experiments). One-way ANOVA with post-hoc Dunnett's test versus WT, $^{*}p < 0.05$.

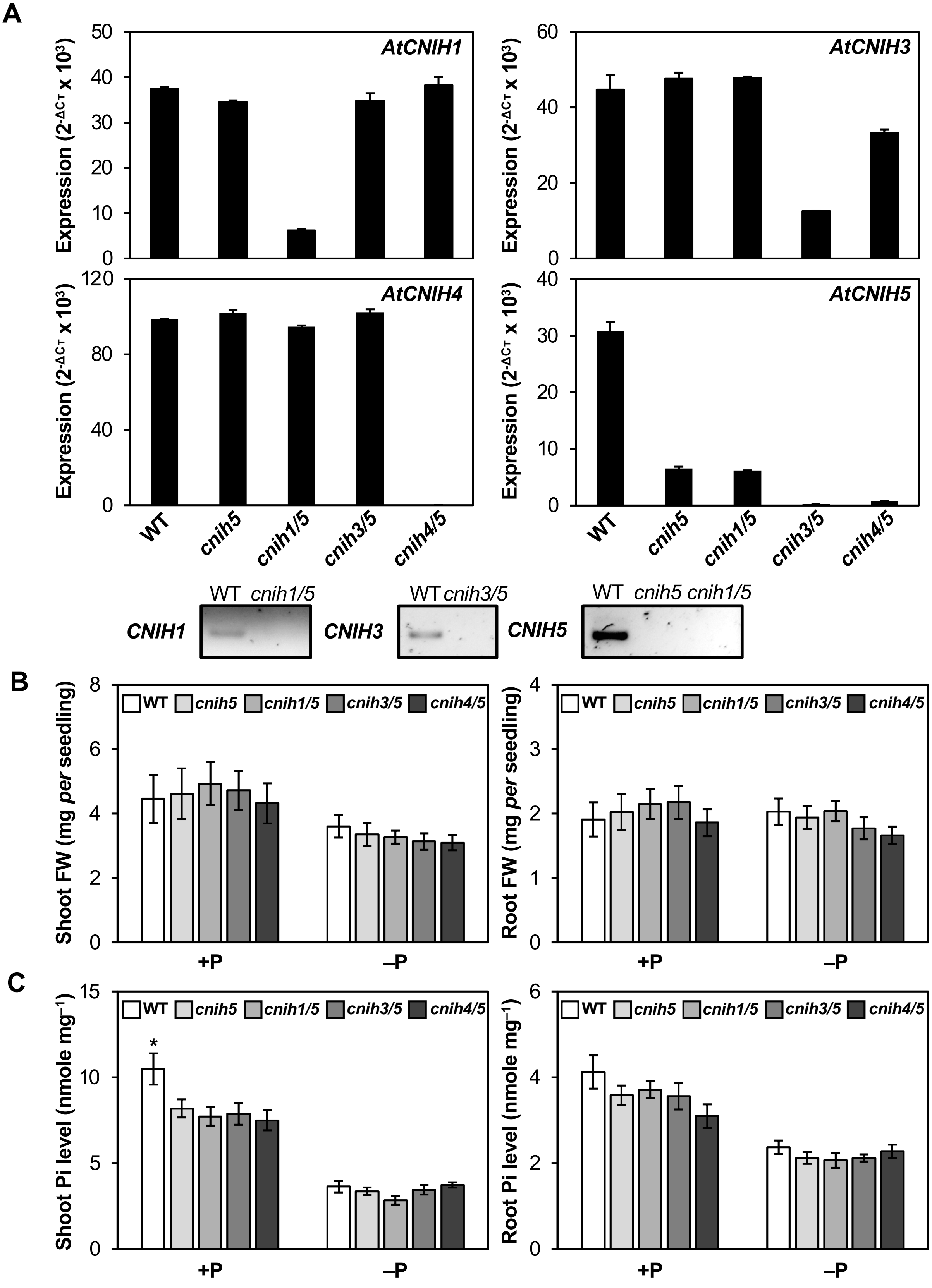


**Fig. S7** Analyses of *Arabidopsis cnih1/5, cnih3/5*, and *cnih4/5* seedlings. (A) The transcript expression of *AtCNIH* genes in the root of 11-day-old *Arabidopsis* WT, *cnih5*, *cnih1/5*, *cnih3/5*, and *cnih4/5* seedlings grown under +P (250 μM $KH_2PO_4$) conditions assessed by RT-qPCR or RT-PCR. (B, C) The shoot and root FW (B) and Pi levels (C) of 11-day-old *Arabidopsis* WT, *cnih5*, *cnih1/5*, *cnih3/5*, and *cnih4/5* seedlings grown under +P (250 μM $KH_2PO_4$) or –P (0 μM $KH_2PO_4$, three days of starvation) conditions. Error bars represent SE (n = 6–10, pools of seedlings from independent experiments). One-way ANOVA with post-hoc Dunnett's test versus *cnih5*, $^*p < 0.05$.

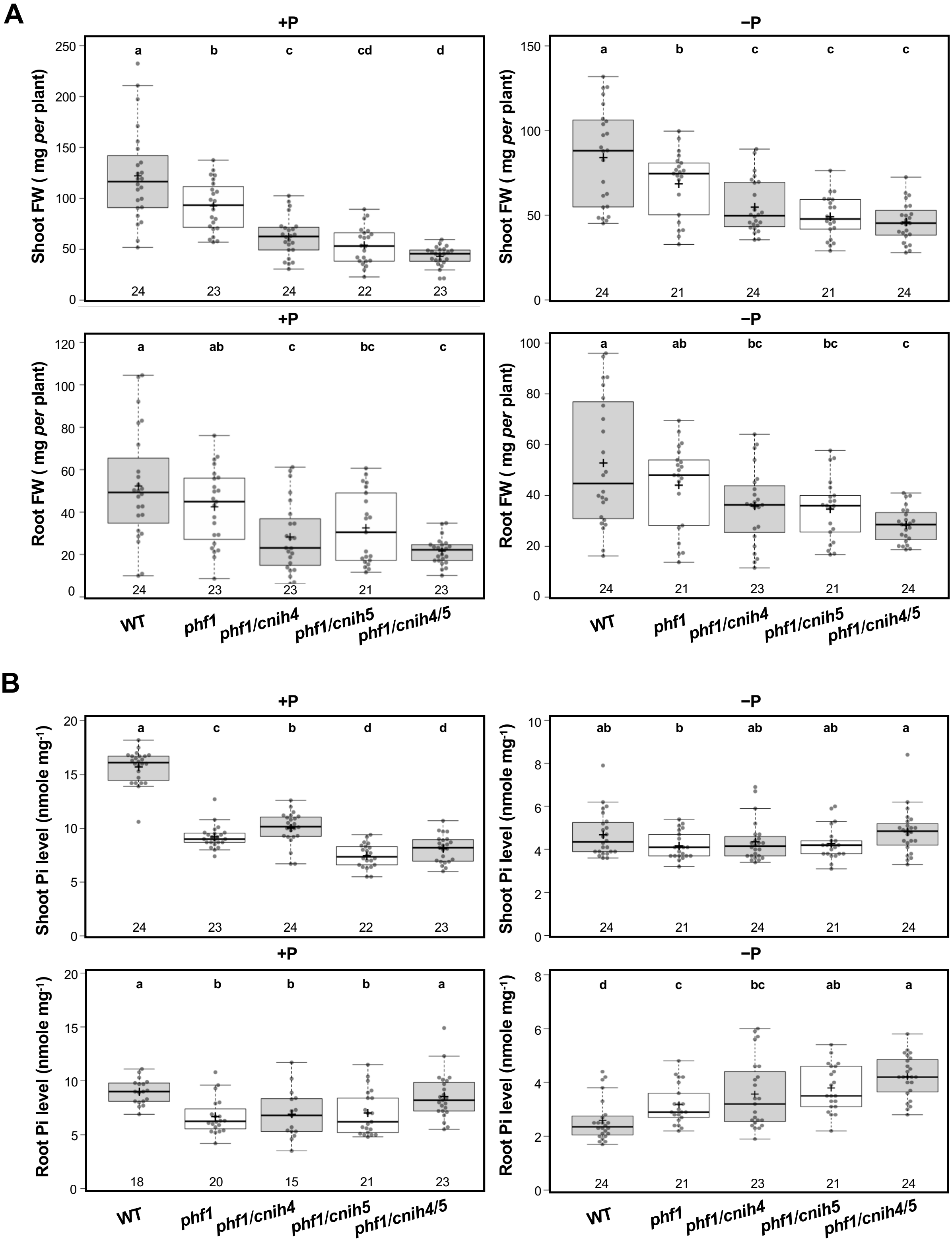


**Fig. S8** Phenotypes of *Arabidopsis phf1, phf1/cnih4, phf1/cnih5*, and *phf1/cnih4/5* adult plants. (A, B) The shoot and root FW (A) and Pi levels (B) of 20-day-old *Arabidopsis* WT, *phf1*, *phf1/cnih*4, *phf1/cnih5*, and *phf1/cnih4/5* plants grown under +P (250 µM $KH_2PO_4$) or –P (10 µM $KH_2PO_4$, five days of starvation) hydroponic conditions. Data are visualized using BoxPlotR (Spitzer *et al*., 2014). Centre lines show the medians; the central plus signs (+) show the means; box limits indicate the 25th and 75th percentiles; whiskers extend to the 5th and 95th percentiles, and data points are plotted as dots. Data significantly different from the other groups are indicated by different lowercase letters (n = 15–24, biological replicates from independent experiments; One-Way ANOVA, Duncan's test).

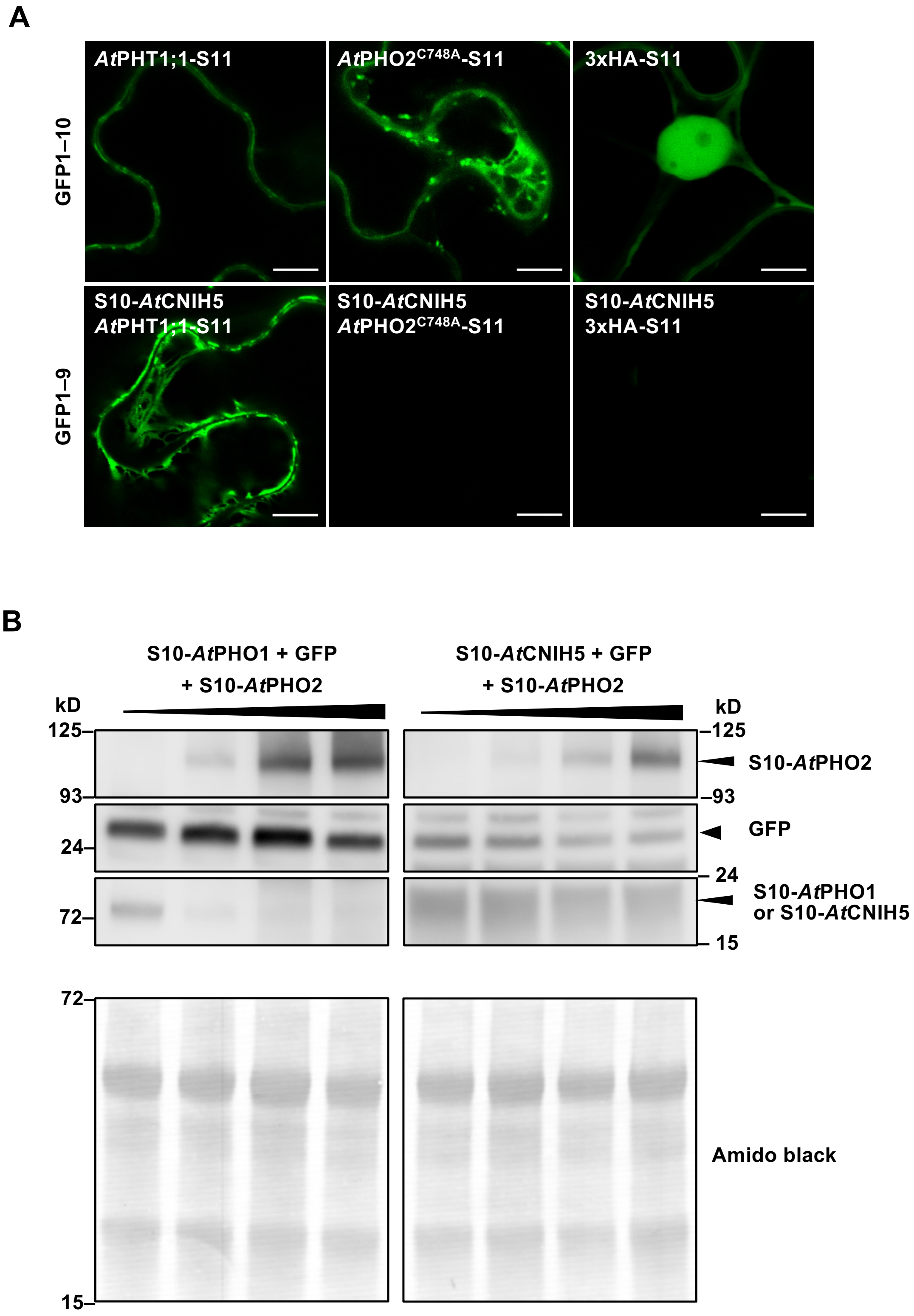


**Fig. S9** *At*PHO2 does not mediate the degradation of *At*CNIH5. (A) Co-expression of S10-*At*CNIH5 with the catalytically defective *At*PHO2$^{C748A}$-S11 in the presence of GFP1–9 in *N. benthamiana* leaves. Images are from single confocal slices. Scale bars, 20 µm. The co-expression of S10-*At*CNIH5 with *At*PHT1;1-S11 and S10-3xHA-S11 are, respectively, used as positive and negative protein–protein interaction controls. (B) Immunoblot analysis of S10-*At*CNIH5 and S10-*At*PHO1 in *N. benthamiana* leaves under co-expression of increasing levels of S10-*At*PHO2. The co-expression of GFP is used as a negative control. Anti-S10 antibodies also detect free GFP.

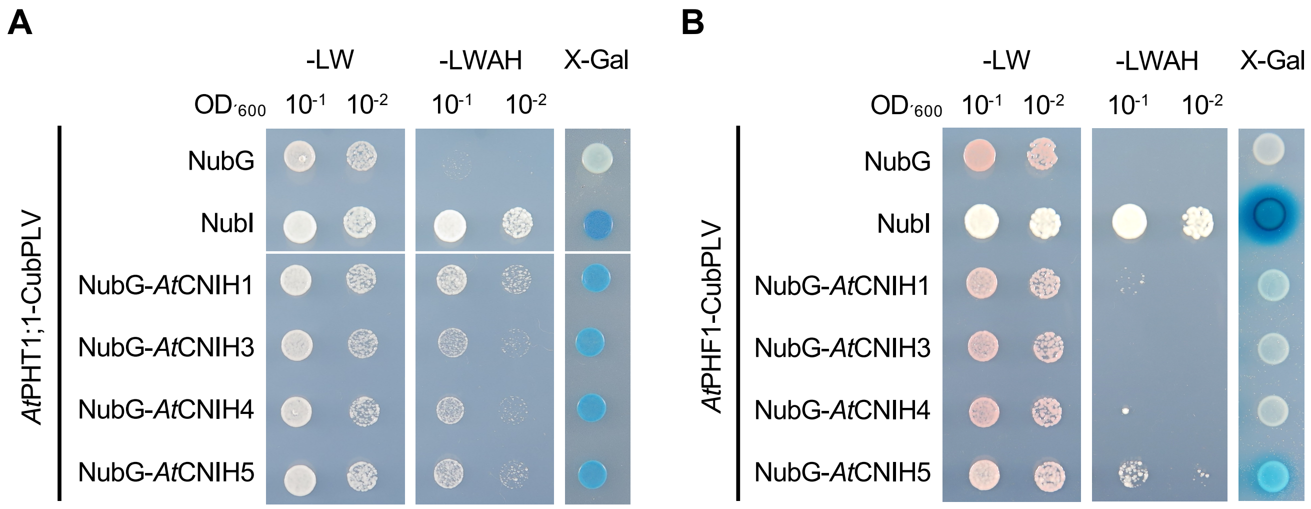


**Fig. S10** Interaction analysis of *At*PHT1;1 and *At*PHF1 with *At*CNIH1, *At*CNIH3, and *At*CNIH4. (a, b) Co-expression of *At*PHT1;1-PLVCub (A) and *At*PHF1-PLVCub (B) with NubG-*At*CNIH1, -*At*CNIH3, and -*At*CNIH4 in the yeast split-ubiquitin system. The co-expression of *At*PHT1;1-PLVCub with NubI and NubG are used as positive and negative controls, respectively. Yeast transformants were grown on synthetic medium without leucine and tryptophan (–LW; the left panel) or on synthetic medium lacking leucine, tryptophan, adenine, and histidine-containing 0.5 μM methionine (–LWAH; middle panel) or on SC–LW containing 2 mg/L X-gal (X-gal; the right panel).

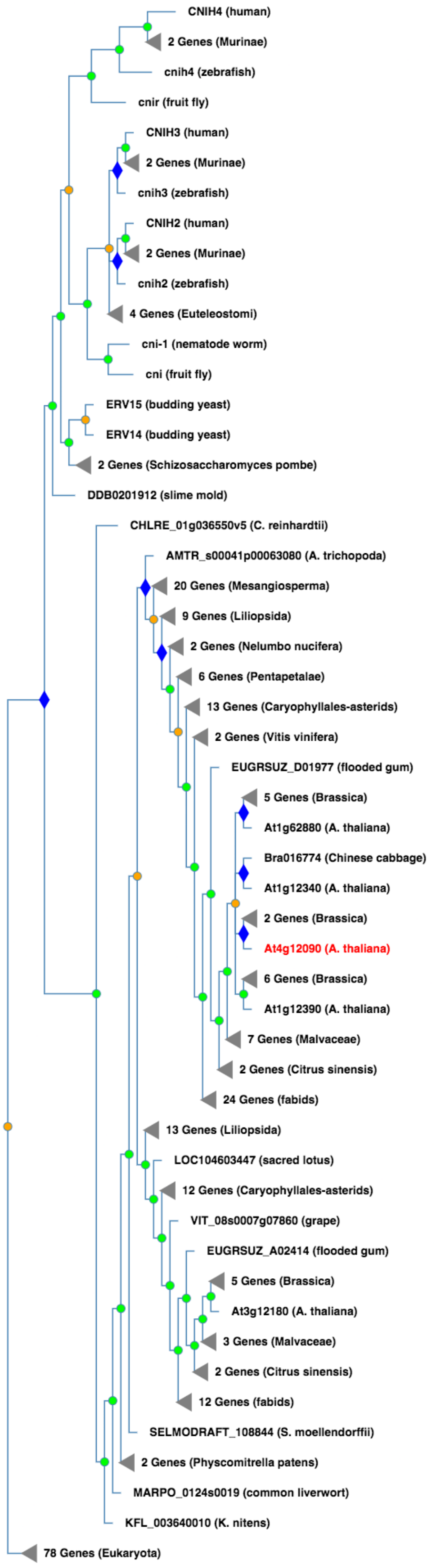


**Fig. S11** The CORNICHON-RELATED family (PTHR12290) in Protein Analysis THrough Evolutionary Relationships (PANTHER). Green circles represent speciation events, while orange circles represent duplication. Diamonds are expanded subfamily nodes, and triangles are collapsed nodes. Note that we have collapsed some nodes (shown with triangles) here to simplify the diagram by hiding some descendant subtrees. Different colors correspond to different subfamilies. The whole family tree can be explored at www.pantherdb.org/treeViewer/treeViewer.jsp?book= PTHR12290

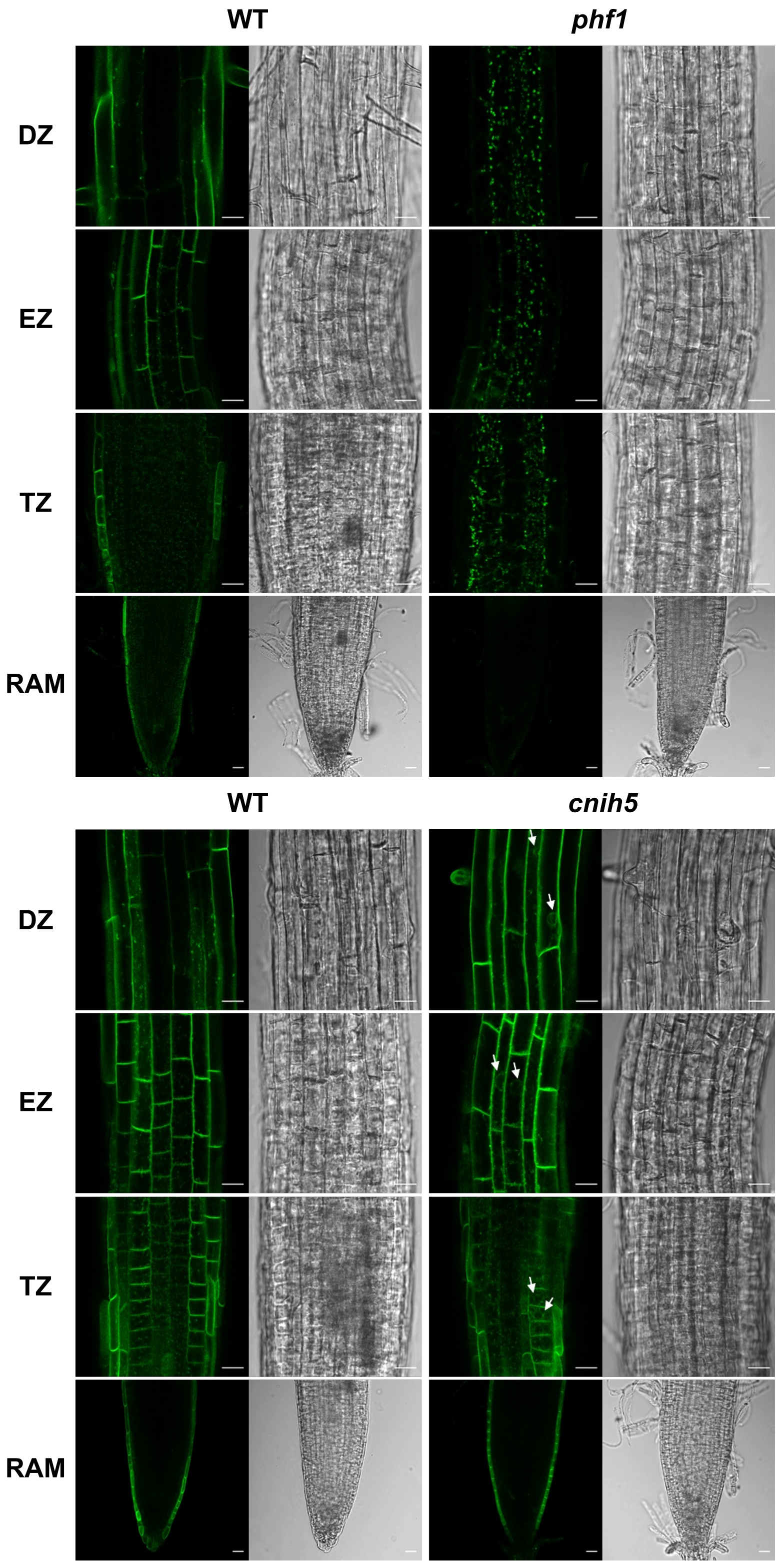


**Fig. S12** Impaired plasma membrane targeting of *At*PHT1;1-S11 in *Arabidopsis phf1*. Representative images of localization of *At*PHT1;1-S11 in 5-day-old *AtPHT1;1pro*: *At*PHT1;1-S11/*35S*: GFP1–10 seedlings grown under −P (0 μM $KH_2PO_4$) conditions. Independent lines in the *phf1* and *cnih5* backgrounds, and the corresponding WT controls are used for comparison. Arrows indicate GFP signals surrounding the nuclear membrane. RAM, Root apical meristem; TZ, transition zone; EZ, elongation zone; DZ, differentiation zone. Images are from single confocal slices. Scale bar, 20 μm.

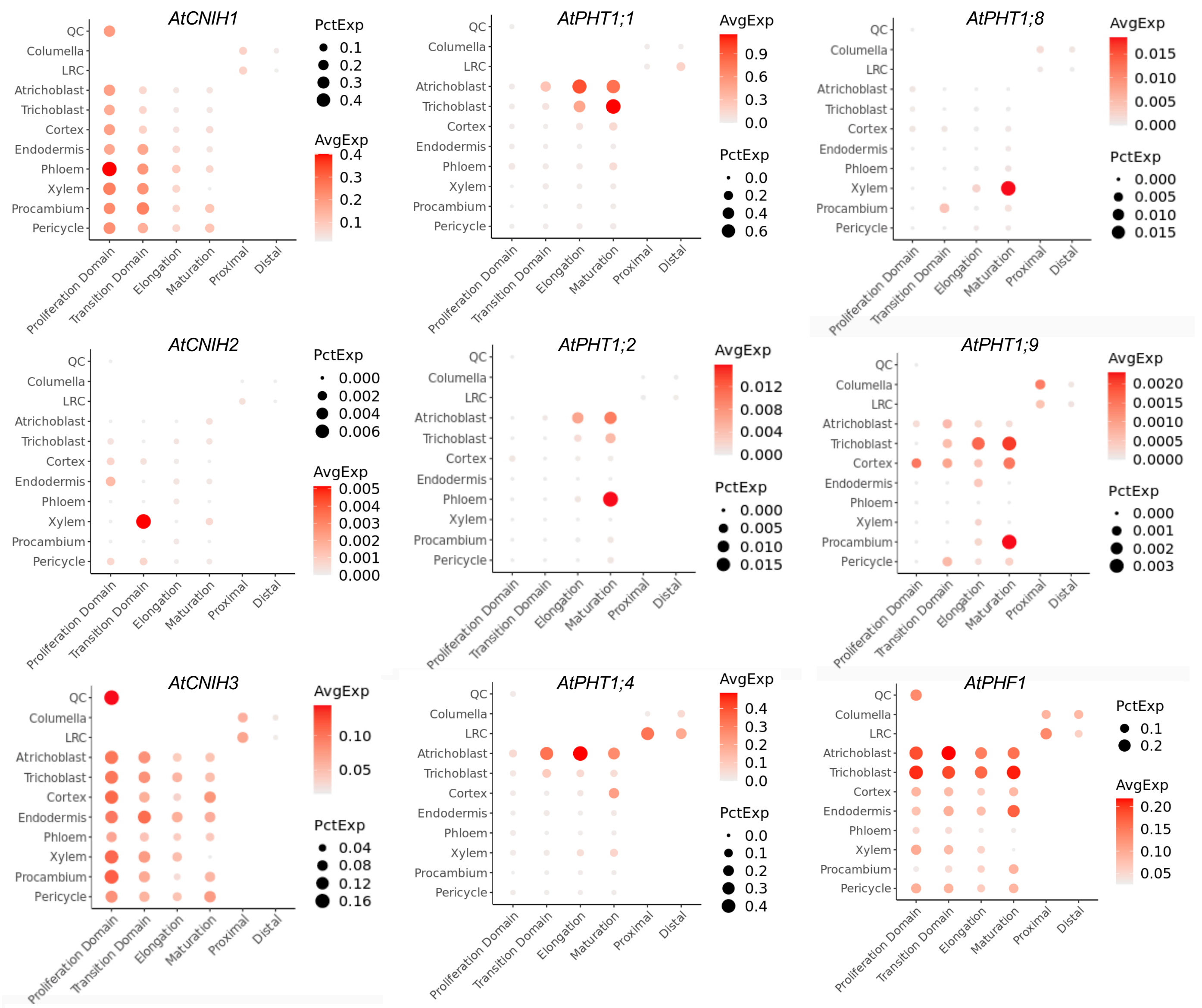


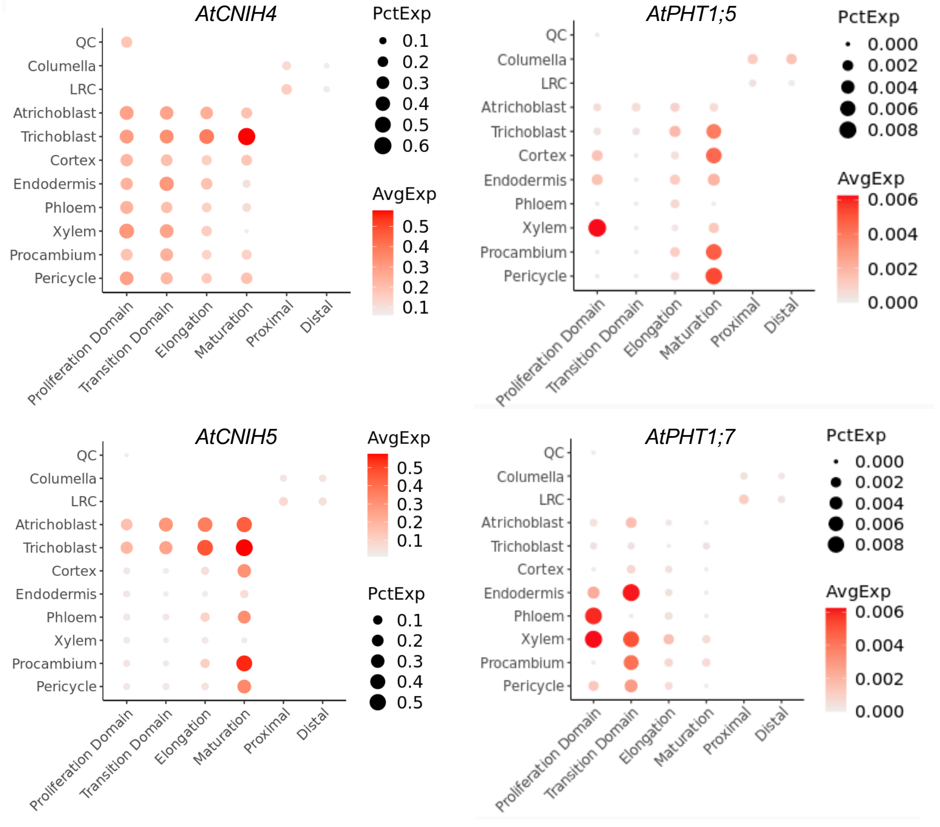


**Fig. S13** Expression of *AtPHT1;1*, *AtPHT1;2*, *AtPHT1;4*, *AtPHT1;5*, *AtPHT1;7–1;9*, *AtCNIH1–5*, and *AtPHF1* revealed by single-cell mRNA sequencing (scRNA-seq) of *Arabidopsis* root. The dot plot used to visualize single-cell gene expression of *Arabidopsis* root is retrieved from ARVEX (https://shiny.mdc-berlin.de/ARVEX/), which hosts processed scRNA-seq data of wild-type samples from Shahan *et al.*, 2022 and Trevor *et al.*, 2023. PctExp: Percentage of cells expressing the gene in a specific time zone and cell type; circle size represents the proportion (%) of cells expressing a given gene. AvgExp: Averaged normalized gene expression value in a specific time zone and cell type; the colour of the scale bar represents the mean expression (natural log +1 pseudocount).

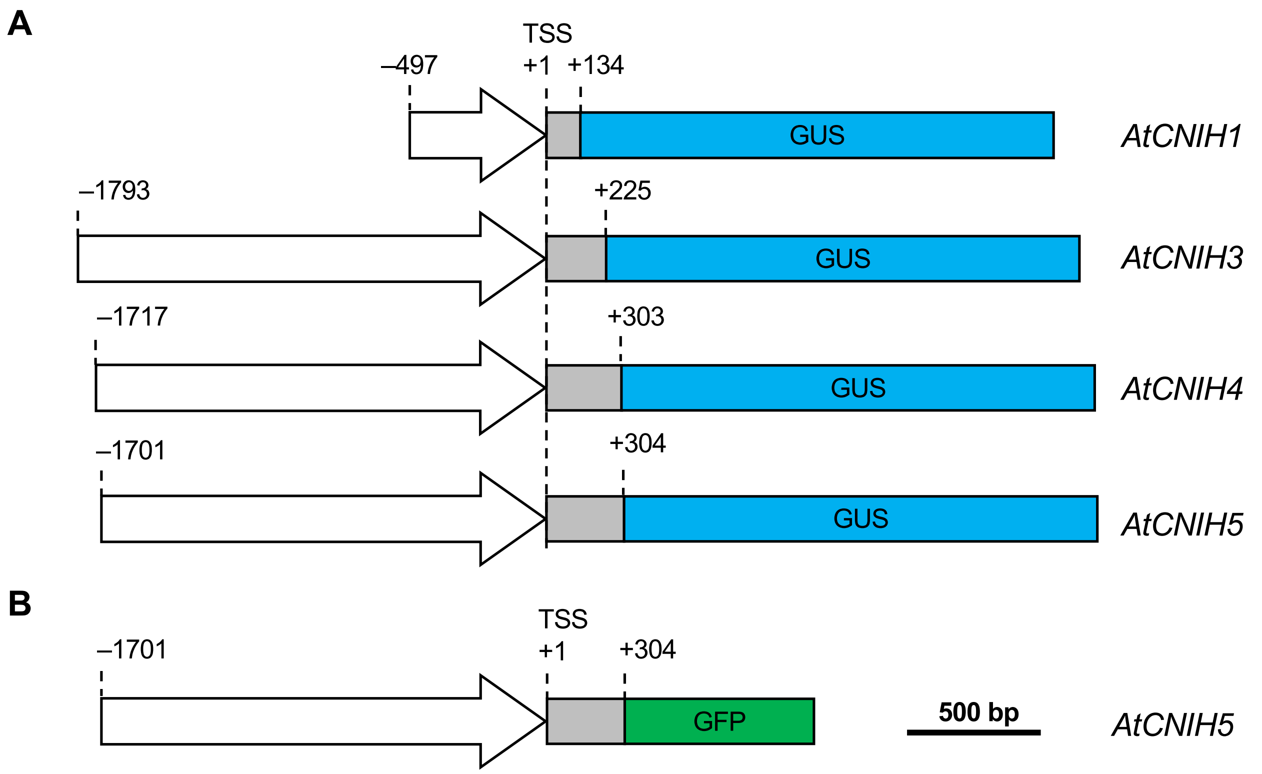


**Fig. S14** Schematic representations of *AtCNIH* gene promoter-fused GUS and GFP reporter constructs. Schematic representations of *AtCNIH1*, *AtCNIH3*, *AtCNIH4*, and *AtCNIH5* promoter-fused GUS reporter constructs (A) and *AtCNIH5* promoter-fused GFP reporter construct (B). The upstream region of transcription started site (TSS) in *AtCNIH* genes was shown by a white arrow, and gray boxes showed the 5’ untranslated regions.

**Table S1. Oligonucleotides used for PCR genotyping, RT-PCR, and RT-qPCR analyses.**

| Gene/AGI No. | Primer name | Sequence (5' to 3') |
|---|---|---|
| **Genotyping** | | |
| *cnih4/AT1G12390* | AscI_*At*CNIH4.for | ggcgcgccaATGGGAGATATCTGGACATGGCTTA |
| | *At*CNIH2,4 STOP_PacI.rev | ttaattaaTCAGTCCTCGTAATCATCCAGCGCCGA |
| | q*At*CNIH4.for2 | AGGAGAGAGAGAGAGAGAGAAGA |
| | SALK_LB1.3.rev | ATTTTGCCGATTTCGGAAC |
| *cnih5/AT4G12090* | AscI_*At*CNIH5.for | ggcgcgccaATGGGCGATCTACTCGATTGGATTA |
| | *At*CNIH5 STOP_XhoI.rev | ctcgagTTAGATGTCTCCTAATGCCGAGTGGA |
| | SAIL_LB1.rev | GCCTTTTCAGAAATGGATAAATAGCCTTGCTTCC |
| | *At*CHIN5(STOP)_BamHI.rev | ggatccTTAGATGTCTCCTAATGCCGAGTG |
| *phf1/AT3G52190* | SALK_037068 LP.rev | CTATGATCCTCAGAAGGCGTG |
| | SALK_037068 RP.for | GTTTTGTGCTGCTCAAAGAGG |
| | SALK_LB1.3.rev | ATTTTGCCGATTTCGGAAC |
| *pho2/AT2G33770* | *PHO2* WT SNP.for | GAAAGTCCAGCAAGAGT**G** |
| | *pho2* SNP.for | GAAAGTCCAGCAAGAGT**A** |
| | *PHO2* WT SNP.rev | AAAGCCCATCGTGATAT |
| **RT-PCR** | | |
| *CNIH1/AT3G12180* | EcoRI_*At*CNIH1.for | gaattcATGGCTTGGGATTTGTTTTTATGGA |
| | *At*CNIH1noSTOP_EcoRI.rev | gaattcAAACAAATGAAGTAGATCATCATCTT |
| *CNIH3/AT1G62880.1* | EcoRI_*At*CNIH3.for | gaattcATGGGAGAGGTCTGGACATG |
| | *At*CNIH3 noSTOP_EcoRI.rev | gaattcGTCCTCGTAATCATCAAGTGTTG |
| *CNIH5/AT4G12090* | EcoRI_*At*CNIH5.for | gaattcATGGGCGATCTACTCGATTGGATTA |
| | *At*CNIH5 STOP_EcoRI.rev | gaattcTTAGATGTCTCCTAATGCCGAGTGG |
| **qRT-PCR** | | |
| *ACT8/AT1G49240* | q*At*ACT8.for | CCCAAAAGCCAACAGAGAGA |
| | q*At*ACT8.rev | CATCACCAGAGTCCAACACAAT |
| *CNIH1/AT3G12180* | q*At*CNIH1.for2 | GTTCCGTGGTATCAGCTTTGA |
| (Fig. 1A) | q*At*CNIH1.rev2 | TATAGACCGCAGAGAGAGTGAG |
| *CNIH1/AT3G12180* | q*At*CNIH1.for3 | GTCGCTTCCGTCTTCTACCAG |
| (Fig. S7A) | q*At*CNIH1.rev3 | GAGTGACCCTTGGAGGATAAAC |
| *CNIH3/AT1G62880.1* | q*At*CNIH3.1.for | CACTTGATGATTACGAGGACTGA |
| (Fig. 1A) | q*At*CNIH3.1.rev | GCGGTGTCAAGTACTAAGAGAG |
| *CNIH3/AT1G62880.1* | q*At*CNIH3.for2 | GAGAGGTCTGGACATGGATTATTT |
| (Fig. S7A) | q*At*CNIH3.rev2 | TTCTTGATGCAGAGTCGTAAGG |

| | | |
|---|---|---|
| *CNIH4/AT1G12390* (Fig. 1A) | q*At*CNIH4.for2 | AGGAGAGAGAGAGAGAGAGAAGA |
| | q*At*CNIH4.rev2 | GCGATGAGGAAGAAGAAGGAAATA |
| *CNIH4/AT1G12390* (Fig. S7A) | q*At*CNIH4.for3 | CTCGAAGCGACAACACTTAG |
| | q*At*CNIH4.rev3 | CATCCAGCGCCGAATAAATC |
| *CNIH5/AT4G12090* (Fig. 1A) | q*At*CNIH5.for | AAAGCAAGGCGGAGATTGA |
| | q*At*CNIH5.rev | AAGCGTTGCGAGGAATAAGA |
| *CNIH5/AT4G12090* (Fig. S7A) | q*At*CNIH5.for3 | CTCTGTTACGTCTGCCAAATG |
| | q*At*CNIH5.rev3 | CACGTCCTCGCCGTCTAA |
| *PHT1;1/AT5G43350* | q*At*PHT1;1.for | GCCATGACGAGAAATAATTATGT |
| | q*At*PHT1;1.rev | TAACTTAAGGTCAACGAGCCAAT |
| *PHT1;2/AT5G43370* | q*At*PHT1;2.for | AGCCATCATTGGAGCCTTC |
| | q*At*PHT1;2.rev | ACCTTAGCCTTGTCTTGATT |
| *PHT1;3/AT5G43360* | q*At*PHT1;3.for1 | CGAGGCTGAGGTTGATAAATGAT |
| | q*At*PHT1;3.rev1 | CACACATCGCAAAACCAATGAC |
| *PHT1;4/AT2G38940* | q*At*PHT1;4 3'UTR.for1 | CGGTCCCAATAGTTTAGGTGATAT |
| | q*At*PHT1;4 3'UTR.rev1 | AGTTGCTAGAGACAAGGAGAA |
| *PHF1/AT3G52190* | q*At*PHF1.for1 | GCCTTCTGAGGATCATAGTAGGTG |
| | q*At*PHF1.rev1 | CGCTGCAACAAGATAAGGAAG |
| *ACP5 /AT3G17790* | q*At*ACP5.for1 | GGTGACGCAGAAGCTCAGCT |
| | q*At*ACP5.rev1 | CCAACTCTGCATCAACGACAA |

**Table S2. Oligonucleotides used for plasmid constructs.**

| Gene | Primer name | Sequence (5' to 3') |
|---|---|---|
| **Cloning for binary vectors** | | |
| S10-*At*CNIH5 | AscI_*At*CNIH5.for | ggcgcgccaATGGGCGATCTACTCGATTGGATTA |
| | *At*CNIH5 STOP_XhoI.rev | ctcgagTTAGATGTCTCCTAATGCCGAGTGGA |
| mCherry-*At*CNIH5 | AscI_*At*CHIN5.for | ggcgcgccATGGGCGATCTACTCGA |
| | *At*CHIN5 STOP_BamHI.rev | ggatccTTAGATGTCTCCTAATGCCGAGTG |
| S10-*At*PHT1;1 | Infusion.for | caaatctaggggatccATGGCCGAACAACAACTAGGAGTGC |
| | Infusion_XhoI.rev | tttagctagactcgagTTATTTCTCGTCATGGCTAACCTCAGCC |
| *At*PHT1;1-S11 | *At*PHT1;1 CDS XbaI.for | tctagaATGGCCGAACAACAACTAG |
| | *At*PHT1;1 CDS noSTOP SpeI.rev | actagtTTTCTCGTCATGGCTAACCTC |
| S10-*At*PHF1 | *At*PHF1_AscI.for | ggcgcgccaATGGAGATTGAAGAAGCGAGTCG |
| | *At*PHF1 STOP_PacI.rev | ttaattaaAGGTCCAAGTTCCACCTACTATG |
| S10-g*At*PHO1 | *At*PHO1_AscI.for | ggcgcgccaATGGTGAAGTTCTCGAAGGAGCTAG |
| | *At*PHO1_PacI.rev | ttaaTTAACCGTCTGAGTCCCTGTCAAGGAACGG |
| S10-*At*PHO2-3xHA | *At*PHO2_AscI.for | ggcgcgccaATGGAAATGTCCCTTACTGACTCTG |
| | *At*PHO2_SpeI no STOP.rev | actagtttTGATTCTGGTCCAATCTCTTGGACGC |
| *At*SEC16A-S11 | AscI_*At*SEC16A.for | ggcgcgccATGGCTTCGACTGCTGA |
| | *At*SEC16A no STOP_SalI.rev | gtcgacCAGTTCAACTTCCTGAAGCTCCTCTCC |
| *At*PHT1;1pro | *At*PHT1;1 pro_ HindIII.for | aagcttATCTCCCAAATGCCGATA |
| | *At*PHT1;1 pro_XhoI.rev | ctcgagACAACGCAAAGAATCCAA |
| *At*PHT1;1-GFP11 | *At*PHT1;1_XbaI.for | tctagaATGGCCGAACAACAACTAG |
| | *At*PHT1;1 no STOP_ SpeI.rev | actagtTTTCTCGTCATGGCTAACCTC |
| *At*CNIH5pro | KpnI_Infusion.for | cctctagaggatccccggGTACCTGAGCATAATCTTCATATGA |
| | Infusion_BglII.rev | CCATagatctCCTCCGCTGTCTTCCGCCGCGTTAC |
| g*At*CNIH5 | BglII_Infusion.for | GGAGGagatctATGGGCGATCTACTCGATTGGATTATCT |
| | Infusion_PacI.rev | gctctagaactagttaattaaCACAAGCTAGCTACCATAACGATAT |
| *At*CNIH1pro | PvuI_*At*CNIH1 pro.for | cgatcgGTTTACCAGTTCAAAAGCTCAGGGTCC |
| | *At*CNIH1 pro_AscI.rev | ggcgcgccATTTGCTCTCTTCAACAATTACGATTCG |
| *At*CNIH3pro | PacI_*At*CNIH3.1 pro.for | ttaattaaGAAGAGAGGAGTAAAAGTAGACAAGAAA |

| | | |
|---|---|---|
| | | CAC |
| | *At*CNIH3.1 pro_AscI.rev | ggcgcgccCCCTCCTCTTCCTCCTGGCTCC |
| *At*CNIH4pro | PacI_*At*CNIH4 pro.for | ttaattaaACCAATCTGTAGCAGTCCAACAAAG |
| | *At*CNIH4 pro_AscI.rev | ggcgcgccCTTCCTCCTTTTGGTTCCGACAATTTC |
| *At*CNIH5pro | PacI_*At*CNIH5 pro.for | ttaattaaCATGCCACTCACAAGTATCGTTGTCA |
| | *At*CNIH5 pro_AscI.rev | ggcgcgccCCTCCGCTGTCTTCCGCCGCGTTAC |
| **Cloning for yeast expression vectors** | | |
| PLVCub-*At*CNIH5 | EcoRI_*At*CNIH5.for | gaattcATGGGCGATCTACTCGATTGGATTA |
| | *At*CNIH5 STOP_EcoRI.rev | gaattcTTAGATGTCTCCTAATGCCGAGTGG |
| NubG-*At*PHT1;4 | MunI_*At*PHT1;4.for | aacaattgATGGCAAGGGAACAATTACAAG |
| | *At*PHT1;4_MunI.rev | aacaattgAACTATTGGGACCGTTCTAC |
